# stGCL: A versatile cross-modality fusion method based on multi-modal graph contrastive learning for spatial transcriptomics

**DOI:** 10.1101/2023.12.10.571025

**Authors:** Na Yu, Daoliang Zhang, Wei Zhang, Zhiping Liu, Xu Qiao, Chuanyuan Wang, Miaoqing Zhao, Baoting Chao, Wei Li, Yang De Marinis, Rui Gao

**Affiliations:** School of Control Science and Engineering, Shandong University, Jinan, Shandong, China; Department of Pathology, Shandong Cancer Hospital and Institute, Shandong First Medical University and Shandong Academy of Medical Sciences, Jinan, Shandong, China; Department of Radiology, Shandong Provincial Hospital Affiliated to Shandong First Medical University, Jinan, Shandong, China; Department of Clinical Sciences, Lund University, Malmö, Sweden; Oxford Centre for Diabetes, Endocrinology & Metabolism, Radcliffe Department of Medicine, University of Oxford, Churchill Hospital, Oxford, U.K

## Abstract

Advances in spatial transcriptomics (ST) technologies have provided unprecedented opportunities to depict transcriptomic and histological landscapes in the spatial context. Multi-modal ST data provide abundant and comprehensive information about cellular status, function, and organization. However, in dealing with the processing and analysis of spatial transcriptomics data, existing algorithms struggle to effectively fuse the multi-modal information contained within ST data. Here, we propose a graph contrastive learning-based cross-modality fusion model named stGCL for accurate and robust integrating gene expression, spatial information as well as histological profiles simultaneously. stGCL adopts a novel histology-based Vision Transformer (H-ViT) method to effectively encode histological features and combines multi-modal graph attention auto-encoder (GATE) with contrastive learning to fuse cross-modality features. In addition, stGCL introduces a pioneering spatial coordinate correcting and registering strategy for tissue slices integration, which can reduce batch effects and identify cross-sectional domains precisely. Compared with state-of-the-art methods on spatial transcriptomics data across platforms and resolutions, stGCL achieves a superior clustering performance and is more robust in unraveling spatial patterns of biological significance. Additionally, stGCL successfully reconstructed three-dimensional (3D) brain tissue structures by integrating vertical and horizontal slices respectively. Application of stGCL in human bronchiolar adenoma (BA) data reveals intratumor spatial heterogeneity and identifies candidate gene biomarkers. In summary, stGCL enables the fusion of various spatial modality data and is a powerful tool for analytical tasks such as spatial domain identification and multi-slice integration.

## Introduction

Cell organization in biological tissues allows interaction between cells and surrounding environment, and spatial information as well as cellular organization on tissue context helps significantly to understand the cellular biological functions and disease pathology^1^. Recent advances in spatial transcriptomics (ST) technologies made it possible to simultaneously capture histological profiles, gene expression profiles and corresponding spatial coordinates that single-cell RNA sequencing (scRNA-seq) cannot capture^2^. The current ST experimental methods can be mainly classified into *in situ* hybridization (ISH) (e.g. seqFISH+^3^, MERFISH^4^) or sequencing-based (ISS) techniques (e.g. STARmap^5^, 10x Xenium^6^) as well as *in situ* capturing-based (ISC) techniques (e.g. Slide-seq^7^, 10x Visium^8^, Stereo-seq^9^). The former measures the spatial distribution of mRNA transcripts with high resolution and accuracy. The latter detects gene expression levels on a genome-wide scale at capture sites (called spots). These emerging ST technologies provide us with new perspectives to study tissue architecture, intercellular interactions, and disease mechanisms.

Analysis leveraging transcriptomics data can elucidate anatomical structures, revealing varied spatial organizations and functionalities within the milieu of complex tissues. For example, conventional ST data clustering methods, such as k-means and Louvain algorithm^10^, typically employ gene expression information to categorize spots into distinct domains. Giotto^11^ and BayesSpace^12^ modeled the spatial structure of neighboring spots using Markov random fields, improving spatial clustering results. SEDR^13^, STAGATE^14^ and GraphST^15^ used deep auto-encoder networks to identify spatially distributed domains by combining expression profiles and spatial locations. Undoubtedly, these methods facilitate the integration of gene expression with spatial cellular distribution, thereby revealing the intricate relationship between spatial gene expression patterns and tissue function across various areas. However, these methods do not sufficiently utilize the multi-modal information in spatial transcriptomics, resulting in downstream analyses that are often characterized by low accuracy and a lack of robustness. K-means and Louvain algorithm only use gene expression information, ignoring spatial information between spots. This can result in discontinuous spatial distributions of spots belonging to the same category in the original spatial context. Methods such as Giotto, SEDR and STAGATE incorporate spatial coordinate information into their algorithms. However, they discard the information on cell organization reflected in histology images. It’s worth noting that some recent proposed algorithms take histology images into consideration. For instance, stLearn^16^ and DeepST^17^ leveraged spatial neighborhood information and morphological features extracted from histology images to normalize gene expression before clustering. SpaGCN^18^ learned color features in the color space of histology images and implemented domain recognition using a graph convolutional network (GCN) that combines gene expression, spatial location, and histology. Despite significant advancements, existing methods exhibit notable limitations. They fall short of effectively capturing high-resolution, content-rich texture features within histology images. Furthermore, these methods have yet to fully tap into their potential to explore and integrate discriminative information from various modalities, thereby impeding their ability to accurately characterize spatial patterns and offer meaningful insights into the biological context.

In addition, spatial transcriptomics sequencing is currently constrained by its inherent technology, restricting its capability to capture tissue samples from a limited region. To study the entire tissue region of interest, the sample should be dissected into multiple slices either vertically or horizontally. That means that ST data typically includes multiple adjacent or consecutive slices of the same tissue. Integrating multiple slices allows for the full utilization of cross-slice information, which yields more comprehensive characterization of spatial organization and potentially enhances the performance of spatial domain identification. At present, only a few computational methods can tackle this integration task. SpaGCN^18^ jointly analyzed multiple serial slices by creating a block adjacency matrix and concatenating gene expression matrices, and aligned multiple adjacent slices by revising the spatial coordinates. However, it ignored the correction of the spatial location between slices. STAGATE^14^ constructed the three-dimensional spatial neighbor network (3D SNN) to integrate multiple consecutive slices for 3D spatial domain identification. Even so, STAGATE required aligned slices as input and cannot handle horizontal integration of multiple adjacent slices. The recently proposed GraphST^15^ utilized the PASTE^19^ algorithm to align spatial coordinates and integrated multiple tissue slices for joint analysis. However, GraphST only considered 2D coordinates, which restricted the accurate depiction of the 3D spatial tissue structure of multiple slices in the real world. The alignment of spatial coordinates across tissue slices remained extremely challenging because of the differences in the structure of the slices and their position on the array^19^. Therefore, the effective and precise integration of spatial data, whether horizontal or vertical, is vital in ST research.

To overcome the above limitations, we develop a versatile cross-modality fusion framework named stGCL based on multi-modal graph contrastive learning for spatial domain detection, multi-slice integration, and the related downstream analyses. stGCL adopts the histology-based Vision Transformer (H-ViT) method, tailored specifically for histology images, to effectively encode histological features such as morphology and spatial distribution of tissue cells. By combining multi-modal graph attention auto-encoder (GATE) with contrastive learning, stGCL deeply fuses transcriptional profiles, histological profiles as well as spatial information simultaneously. By implementing the above two key steps of data processing and fusion, stGCL can generate effective embeddings for accurately identifying spatially coherent regions across transcriptional and histological profiles. In addition, we introduce a multi-slice alignment algorithm based on tissue slice edge structure that ensures efficient integrated analysis of ST data across slices, uncovering spatial heterogeneity in 3D tissue structure on a broader scale. We conducted extensive experimental tests on spatial transcriptomics data obtained from various experimental platforms, elucidating the superior performance of stGCL from multiple perspectives including domain identification accuracy, multi-slice integration, robustness, and biological mechanism analysis. As a proof-of-concept, we also applied stGCL to in-house bronchiolar adenoma (BA) distinguishing tissue spatial heterogeneity and providing insights into the underlying mechanisms of the tumor.

## Results

### Overview of stGCL

The overall framework of stGCL is depicted in **Fig. 1A**. In brief, the core algorithm of stGCL takes gene expression, spatial location, as well as histology image as input and outputs the spot embedding reflecting expression similarity, morphological similarity, and spatial proximity, which can be further used for exploring spatial domains in differential scenarios. Certainly, stGCL possesses the capability to effectively integrate multi-slice data and the latent space representation can be directly utilized for other tasks, such as spatial domain identification, low-dimensional visualization, trajectory inference and cell-cell communication analysis (**Fig. 1D**).

**Fig. 1.**
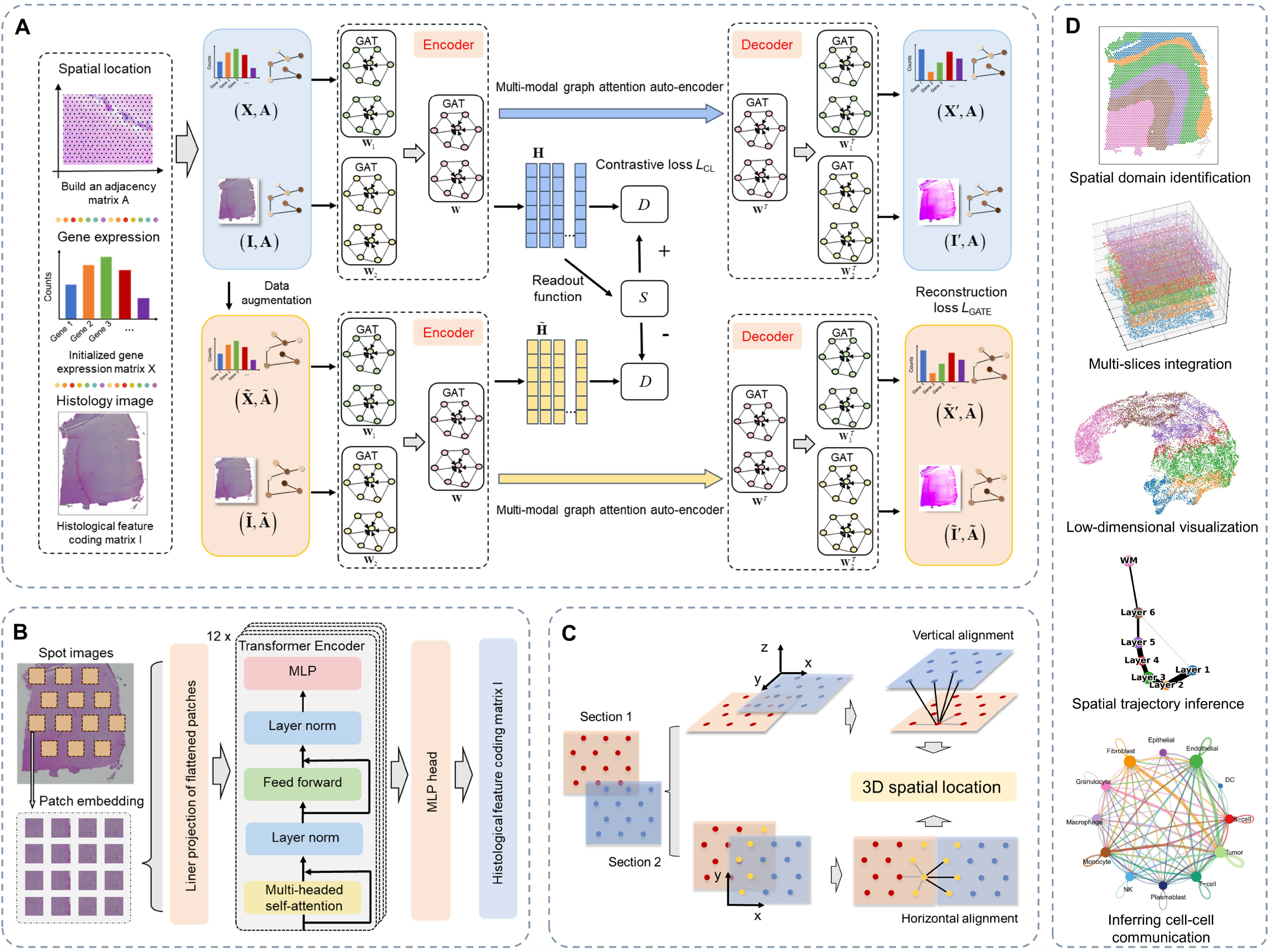
Schematic diagram of the stGCL. **A**. stGCL integrates gene expression profiles, spatial information and histological profiles as input, subsequently deriving low-dimensional latent representations through multi-modal graph contrastive learning. In particular, stGCL aims to minimize reconstruction loss and contrastive loss during the training phase. This process involves the utilization of multi-modal GATE and contrastive learning techniques to produce a comprehensive spot joint embedding. **B**. stGCL utilizes a modified H-ViT model for the extraction of morphological features from histology images. **C**. In ST datasets with multiple tissue slices, stGCL vertically or horizontally aligns these sections along the edge of the cut surface to maintain spatial consistency. After alignment, it constructs a spatial neighborhood graph using the coordinated locations, encompassing location data from all slices. **D**. The output of stGCL is adaptable for diverse downstream tasks, including spatial domain identification, trajectory inference, and cell-cell communication inference.

The stGCL adopts modified H-ViT to efficiently encode the histological features of the spots by dividing each spot image into multiple patches, thereby obtaining the histological feature encoding matrix **I** (**Fig. 1B**). Then a spatial neighborhood graph is constructed based on spatial coordinates to characterize the relationship between the spots. Next, stGCL uses data augmentation, multi-modal GATE and contrastive learning to embed spot gene expression, histological information and spatial similarity for generating the low-dimension embedding **H** (**Fig. 1A**). Specifically, the proposed multi-modal GATE learns spot embedding by iteratively aggregating gene expression features and histological features from adjacent spots. This step encodes each spot and its corresponding negative sample (see Methods), and then decode the embeddings to reconstruct gene expression and histological profiles. In contrastive learning, stGCL maximizes the mutual information between the node representation and the global information of the entire graph, thereby endowing the spot embedding with both local and global structural patterns. Finally, stGCL combines reconstruction loss and contrastive loss to update the spot embedding **H** to make it more informative and discriminative.

In addition to processing individual slices, stGCL also possesses embedding capabilities for horizontal and vertical multi-slice data (**Fig. 1C**). The spatial coordinates of multiple slices are first aligned vertically or horizontally using the novel alignment strategy, and a spatial neighbor network is built (see Methods). Then stGCL integrates gene expression information, histological information and spatial information of multiple slices to learn spot embedding with more comprehensive cross-slice information, which encourages smoothing of adjacent spot features within and across slices as well as mitigates the batch effect. Then, spot embedding can be applied for downstream analytical tasks.

### stGCL accurately dissects tissue structures from the dorsolateral prefrontal cortex dataset

To evaluate the performance of stGCL, we applied stGCL to 12 human dorsolateral prefrontal cortex (DLPFC) slices generated by the 10x Visium platform^20^ for spatial domain detection task. Based on cytoarchitecture and gene markers, the DLPFC dataset was manually annotated into neuronal layers (Layer 1-Layer 6) and white matter (WM) (**Fig. 2A**), which serves as ground truth. For performance comparison, we benchmarked stGCL against other methods, including the non-spatial clustering approach (SCANPY^21^) as well as six recently proposed spatial clustering methods (BayesSpace^12^, SpaGCN^18^, SEDR^13^, STAGATE^14^, DeepST^17^, and GraphST^15^). These methods were applied to the DLPFC dataset using their recommended default parameters, allowing for a comprehensive and fair assessment of stGCL’s capabilities in relation to current methodologies.

**Fig. 2.**
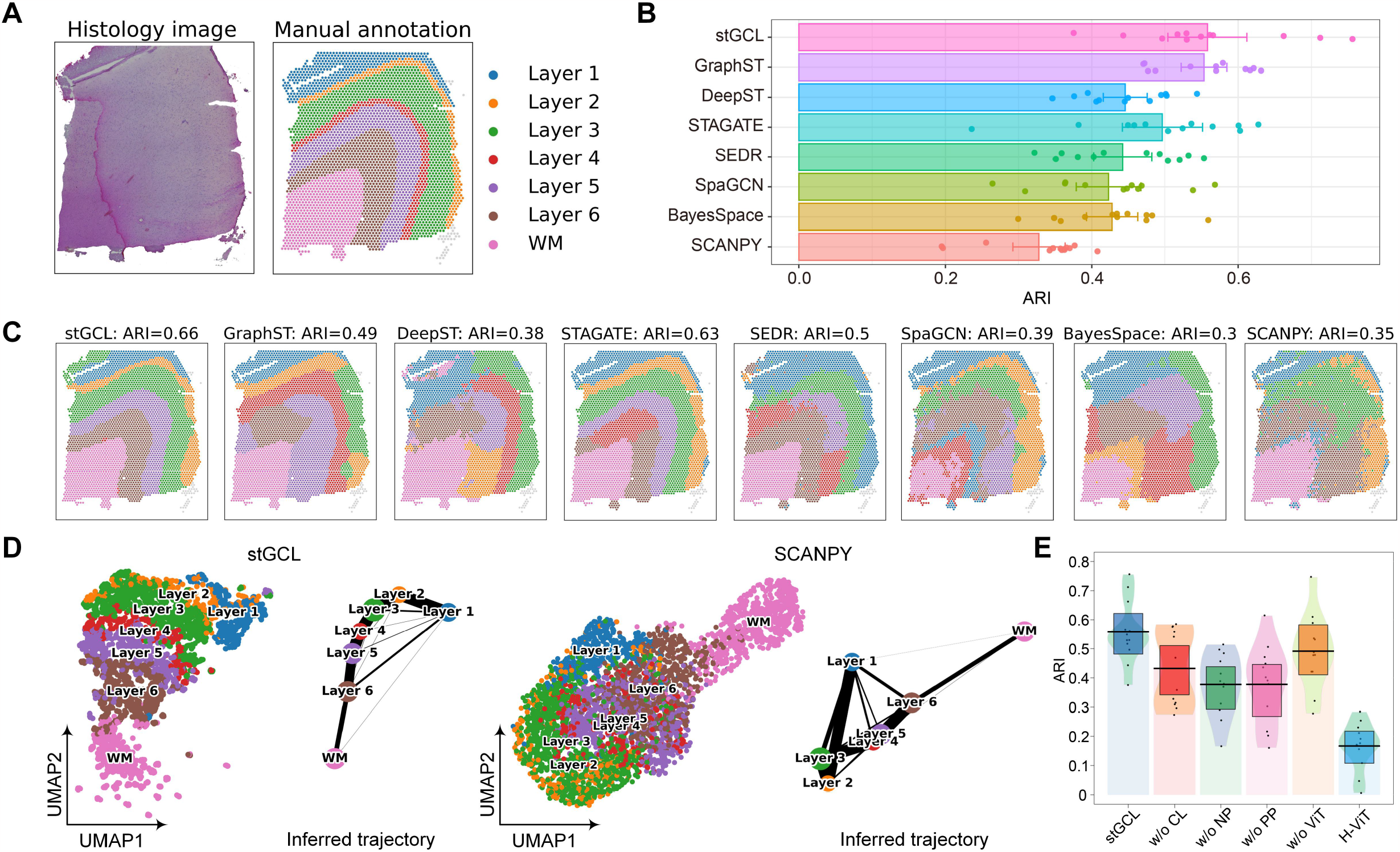
stGCL enables accurate identification of layer structures in the DLPFC data. **A**. Hematoxylin and eosin (H&E) staining image and manually annotated tissue structures for DLPFC slice 151674. **B**. Clustering results of eight methods on 12 DLPFC slices. The results are evaluated by ARI scores. **C**. Spatial domains identified by eight methods within the DLPFC slice 151674. **D**. UMAP visualizations and PAGA graphs for DLPFC slice 151674 from the embeddings of stGCL and SCANPY. **E**. The ARI pirate graph of ablation studies for stGCL. The ARI scores of each slice is represented by points. Horizontal lines represent the mean scores for each condition and the boxes denote the 95% confidence intervals.

In the analysis of 12 slices, stGCL demonstrated superior efficiency in identifying brain tissue structures, achieving the highest level of clustering performance (mean ARI = 0.56) (**Fig. 2B and Supplementary Fig. S1**). It is noteworthy that clustering methods based on spatial information significantly outperform those not utilizing spatial data (**Fig. 2B, C**). This indicates that considering spatial neighborhood information is crucial for the accurate identification of functional regions in tissues. Additionally, we noted that despite both SpaGCN and DeepST incorporating histological information, their performance falls short compared to SEDR, STAGATE, GraphST, and stGCL. This phenomenon is likely attributable to the suboptimal integration strategies of histological information adopted by these methods. Differing from other approaches, stGCL treats histology images as an independent data modality and employs a contrastive learning method for data fusion. This fusion technique enables the full utilization of information from multiple modalities, which is fundamental to stGCL’s superior performance compared to other methods. For instance, in the analysis of slice 151674, which comprises 3,673 spots and 19,856 genes, the clustering results obtained by stGCL exhibits the most clear and continuous cortical layer boundaries (ARI=0.66). Simultaneously, stGCL and STAGATE are the only methods that can accurately depict Layer 2 (**Fig. 2C**). However, stGCL identifies Layer 2 as a thin layer, matching most closely with the shape of the annotated cortical layer.

To better demonstrate the effectiveness of the latent information discovered by stGCL, we conducted further developmental trajectory inference analysis on slice 151674 (**Fig. 2D**). In brief, utilizing the latent embedding information obtained from stGCL, we employ uniform manifold approximation and projection (UMAP) to reveal distances between spatial domains and perform trajectory inference using the partition-based graph abstraction (PAGA)^22^ algorithm. From **Fig. 2D**, it is evident that the individual cortical layers are well organized in the UMAP plot of stGCL. Additionally, we utilized results from SCANPY to plot a corresponding UMAP diagram for comparison. In the SCANPY UMAP plot, only the Layer 1 and white matter (WM) show effective separation, while spots of other layers exhibit a mixed phenomenon. In contrast, the PAGA results derived from the latent information of stGCL distinctly display different layers. Moreover, the PAGA results show a near-linear developmental trajectory from Layer 1 to Layer 6, with greater similarity observed between adjacent cortical layers.

To further test the effectiveness of the core modules of stGCL, ablation experiments were conducted on 12 DLPFC slices. From **Fig. 2E**, we observed that the performance of stGCL decreases by 6.7% when histological information is disregarded (w/o H-ViT). The mean ARI score for encoding histological image features using H-ViT alone is 0.17. This demonstrates that the H-ViT module, which considers histological images, effectively extracts histological information and is undoubtedly necessary for enhancing the performance in domain identification. Additionally, we found that the performance of stGCL is affected by factors such as the absence of contrastive learning (CL), positive and negative pairs (PP and NP) in CL. When stGCL utilizes all modules and achieves feature fusion, it is capable of realizing optimal performance.

### stGCL improves integration of multiple tissue slices

The study of gene expression in relation to spatial context often necessitates the integration of multiple tissue slices to construct a more detailed and comprehensive gene expression atlas. To conduct spatial transcriptomic studies on tissue regions of interest, experimental samples often need to be divided into multiple slices for sequencing, both horizontally and vertically. Consequently, to obtain more complete and continuous spatial transcriptomic information of tissues or organs, computational methods for horizontal or vertical stitching and alignment are now essential^15^. However, most current analysis methods are only applicable to individual tissue slices and cannot jointly identify spatial domains from multiple slices, either horizontally or vertically. Furthermore, batch correction methods developed for scRNA-seq only consider gene expression relationships and neglect spatial information, making batch effects difficult to resolve. Therefore, joint modeling and analysis of multiple slices represent a significant challenge in the field of spatial transcriptomics.

stGCL employs a novel multi-slice alignment algorithm based on tissue slice edge structure to address this challenge. It explores the neighborhood information within each slice and between adjacent slices across batches through spatial alignment of multiple tissue slices. To validate the efficacy of stGCL, we applied it to four consecutive slices from the aforementioned DLPFC dataset (151673, 151674, 151675, and 151676). The first two slices and the last two slices are directly adjacent, and the middle two slices are 300 μm apart^20^. stGCL successfully aligns the four slices in 3D space (**Fig. 3A**). In **Fig. 3C**, it is evident that SCANPY and SEDR mix the four slices together, failing to distinctly differentiate the cortical layers. While STAGATE employs a 3D spatial neighbor network (3D SNN), its results still exhibit batch-specific separations of specific slices. stGCL effectively performs low-dimensional embedding of the multi-slices, producing clear and ordered separations between the different layers (from Layer 1 to Layer 6 and WM). This demonstrates that stGCL not only corrects for batch effects but also preserves tissue heterogeneity information. The outstanding performance of stGCL can be attributed to the fact that the cross-slice spatial neighborhood graph alleviates the feature distribution differences between batches and the spot representation learned by CL covers the global features of tissue slices. Moreover, stGCL achieved the highest ARI score (ARI = 0.61) for the integration of the four slices (**Fig. 3B**). Notably, the integrated analysis using four tissue slices with stGCL demonstrated higher clustering accuracy than the analysis using only one slice (**Fig. 3D and Supplementary Fig. S1**). For instance, stGCL not only captured smooth domain boundaries but also depicted the white matter (WM) structure more closely aligned with the expected truth. This indicates that integrating multiple tissue slices facilitates the consideration of cross-slice information, thereby enhancing the performance of spatial domain identification. Additionally, we conducted a vertical alignment analysis of 12 DLPFC slices using stGCL. Here again, stGCL obtained the highest ARI scores and exhibited superior integration performance (**Supplementary Fig. S2 A-C**).

**Fig. 3.**
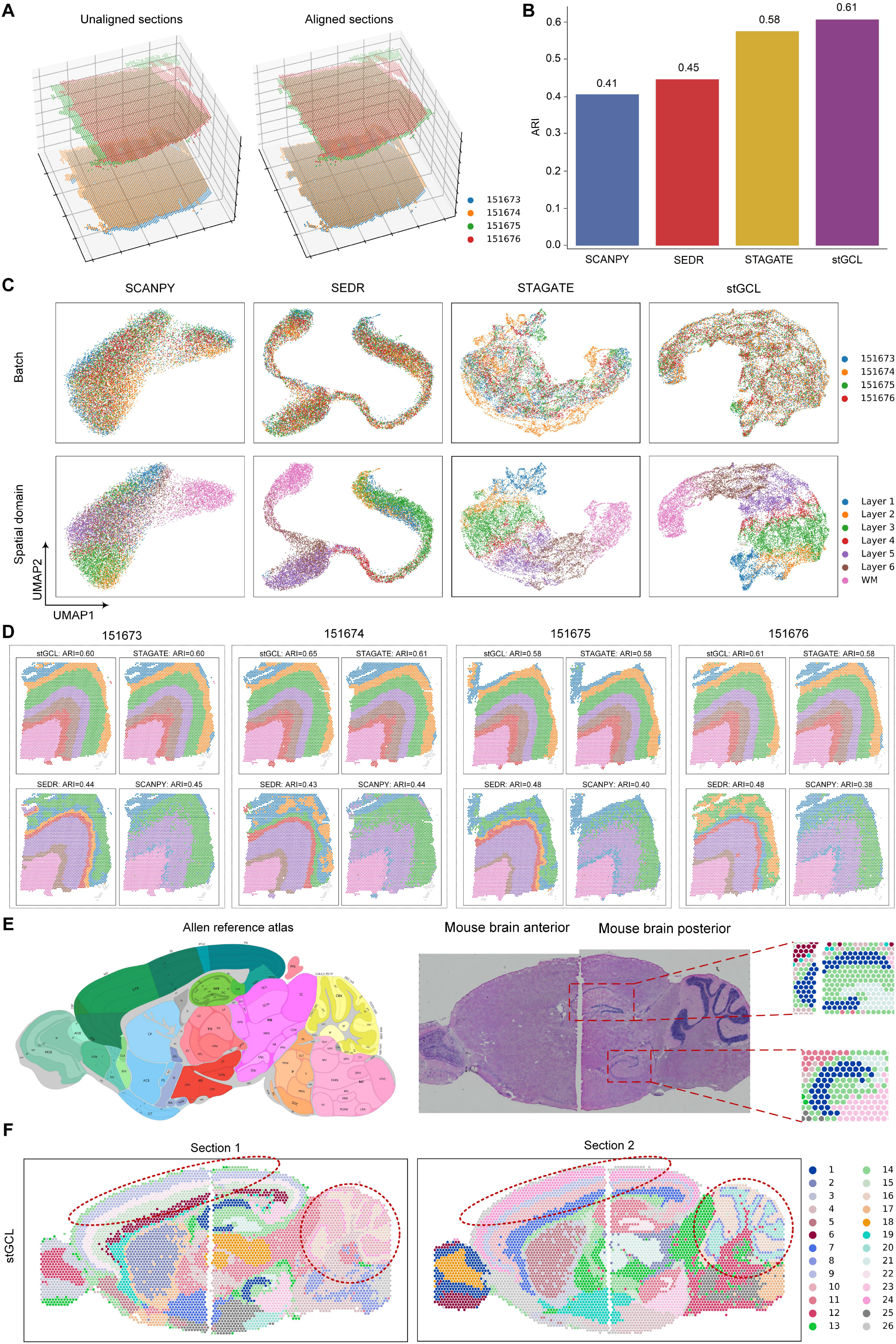
stGCL improves both vertical and horizontal integration within DLPFC datasets, as well as in anterior and posterior sagittal slices of the mouse brain. **A**. Visualization of four DLPFC slices in 3D space (left). The aligned results from stGCL are shown in right panel. **B**. A histogram depicting the results of spatial clustering for four DLPFC slices is shown, executed through four different methods. **C**. The UMAP visualization showcases an integrated analysis of four slices utilizing SCANPY, SEDR, STAGATE, and stGCL. In this representation, spots are color-coded to represent different slices (top) and cortical layers (bottom), respectively. **D**. Identification of spatial domains in four tissue slices using four different integration algorithms. **E**. Anatomical annotations from the Allen Mouse Brain Atlas (left) and H&E images of the mouse brain sagittal anterior and posterior (right). The red boxes indicate the dorsal and ventral horns of the hippocampus region identified by stGCL in section 1, which both contain the cornu ammonis (CA) and dentate gyrus (DG) structures. **F**. Results of horizontal integration of stGCL on two mouse brain sections. Red circles mark the cerebral cortex (CTX) and cerebellum (CBX) regions identified by stGCL, respectively.

Next, we applied stGCL to sagittal anterior slices (section 1 and section 2) of mouse brain^23^, which exhibited evident batch effects. We vertically aligned section 1 and section 2 in 3D space (**Supplementary Fig. S2F**) and the UMAP plots were then used to visually evaluate the integration results (**Supplementary Fig. S2G**). stGCL successfully integrated these two slices, achieving an effective low-dimensional spatial representation (**Supplementary Fig. S2F, G**). This demonstrates stGCL’s ability to efficiently integrate tissue slices from multiple batches and accurately align tissue structures across different slices.

In response to the issue of limited size in spatial transcriptomics capture regions, we also tested the horizontal multi-slice integration capability of stGCL. Here, we adopted two sections of mouse brain data^23^, where both sections were divided into sagittal anterior and posterior slices. For a fair comparison, the number of clusters in stGCL is set to 26, the same with the previous studies SpaGCN^18^ and GraphST^15^. Compared to SpaGCN, the tissue structures represented by stGCL and GraphST align more consistent with annotations in the Allen Reference Atlas^24^ (**Fig. 3E**), while SpaGCN lacks clear spatial separation and exhibits many noisy spots (**Fig. 3F and Supplementary Fig. S3**). stGCL and GraphST are aligned along the shared edges of the anterior and posterior brain, which better reflects the spatial adjacency of the two slices. Furthermore, stGCL outperforms GraphST in capturing complex and fine tissue structures. Specifically, the cerebral cortex (CTX) and cerebellum (CBX) regions delineated by stGCL are more consistent with annotations in the anatomy. In addition, stGCL clearly localizes the dorsal and ventral horns of the hippocampus region, and accurately captures the “cord-like” and “arrow-like” structures (cornu ammonis (CA) and dentate gyrus (DG) domains) within them (**Fig. 3E, F**). Overall, stGCL enables efficient identification of shared layer structures across tissue slices through horizontal integration.

We further explored the biological functionality of the spatial domains identified by stGCL by conducting research on the identified marker genes (**Supplementary Fig. S4**). Neurod6, a gene encoding a neural transcription factor, is highly expressed in the CA region of the hippocampus^25^. C1ql2, a known biomarker for the DG, is upregulated in the DG region^26^. Gng4 shows differential expression in the granule layer of the main olfactory bulb (MOBgr)^27,28^. Prkcd and Lamp5 are respectively enriched in the thalamus and superficial cortex layer^29,30^. These gene expression variations, previously confirmed in studies, have been further validated in the results of stGCL (**Supplementary Fig. S4**). Therefore, these findings underscore the significance of stGCL in accurately identifying domains across tissue slices and successfully revealing spatially specific expression patterns.

### stGCL demonstrates robustness and scalability on spatial transcriptomic data across platforms and resolutions

The analysis algorithms for spatial transcriptomics data exhibit a certain sensitivity to different experimental platforms^31^. This sensitivity is primarily due to variations in sample processing, data acquisition methods, and the quality and characteristics of data across various platforms. For instance, different platforms may differ in terms of spatial resolution, gene detection sensitivity, and the complexity of sample processing. To test the universality of stGCL, we conducted an analysis and comparison on spatial transcriptomics data obtained from different platforms with varying spatial resolutions and spatial modalities.

We first applied stGCL to a human non-small cell lung cancer (NSCLC) dataset captured by NanoString CosMx SMI (high-plex spatial molecular imaging)^32^. This dataset comprises expression profiles and histological profiles with subcellular resolution, containing 20 field of views (FOVs) and 12 cell types (**Fig. 4A**). In comparison, stGCL portrays cell distribution with higher accuracy and distinctly differentiated the dominant tumor cells and fibroblasts, showing consistency that aligns more closely with manual annotation (**Fig. 4A-C, Supplementary Fig. S5A, C**). Subsequently, we utilized the labels obtained from stGCL as grouping criteria and employed CellChat^33^ for the discovery of complex ligand-receptor (L-R) interactions between different cell types in NSCLC (**Supplementary Fig. S5D-G**). The results from stGCL corroborate the cell types that frequently communicate in NSCLC, specifically the strong interactions between fibroblasts, endothelial cells, macrophages, T-cells, and tumor cells, as well as within tumor cells themselves (**Supplementary Fig. S5F, G**)^34^. We further visualized the significant L-R pairs involved in the communication between different cell types (**Supplementary Fig. S5F, G**). The results demonstrate that the L-R pair amphiregulin (AREG) and epidermal growth factor receptor (EGFR) expressed within tumors exhibits the strong interaction. Additionally, as previous research has demonstrated, AREG activates the EGFR signaling pathway, which in turn promotes the growth, proliferation, and metastasis of tumor cells^35^.

**Fig. 4.**
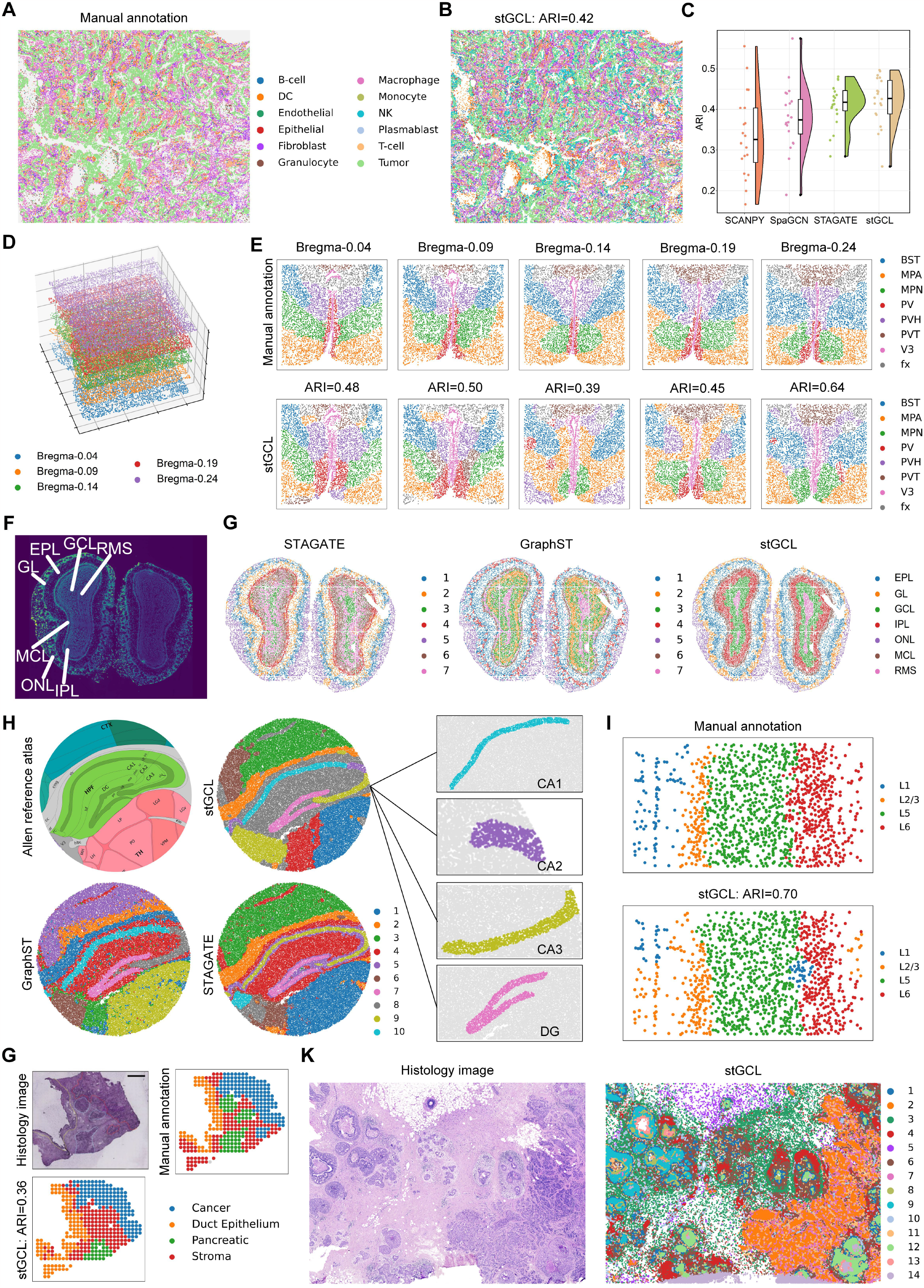
stGCL achieves superior clustering performance on datasets profiled by different ST platforms. **A**. Manual annotation of human NSCLC NanoString CosMx SMI data containing 20 FOVs. **B**. Visualization of the clustering results of stGCL. **C**. Raincloud plot of ARI scores for four methods at 20 FOVs of NSCLC data. **D**. stGCL aligns five consecutive hypothalamic preoptic area slices (Bregma-0.04, Bregma-0.09, Bregma-0.14, Bregma-0.19, and Bregma-0.24) generated by MERFISH in 3D space. **E**. Tissue domain annotations from the previous study (top) and cluster assignments produced by stGCL (bottom). BST: bed nuclei of the strata terminalis; MPA: medial preoptic area; MPN: medial preoptic nucleus; PV: periventricular hypothalamic nucleus; PVH: paraventricular hypothalamic nucleus; PVT: paraventricular nucleus of the thalamus; V3: the third ventricle; and fx: columns of the fornix. **F**. Laminar organization of mouse olfactory bulb Stereo-seq data annotated in DAPI-stained image. RMS: rostral migratory stream; ONL: olfactory nerve layer; IPL: internal plexiform layer; GL: glomerular layer; MCL: mitral cell layer; GCL: granule cell layer; EPL: external plexiform layer. **G**. Clustering results of STAGATE, GraphST and stGCL. **H**. Structural annotation of mouse hippocampus from the Allen Reference Atlas (left) and spatial domains detected by STAGATE, GraphST, and stGCL on mouse hippocampus Slide-seqV2 data (right). The black boxes show the CA1, CA2, CA3, and DG structures identified by stGCL. **I**. Layer structure of mouse mPFC STARmap data and Visualization of the spatial domain identified by stGCL. **G**. Histology image of PDAC with manual annotations (cancer, duct epithelium, pancreatic and stroma); Clustering results using stGCL embeddings on PDAC data profiled by ST technology. **K**. H&E stained image and spatial domains detected by stGCL on breast cancer 10x Xenium data.

We further evaluated the performance of stGCL under the MERFISH technology which offers single-cell level resolution but detects less number of genes^36^. For the mouse hypothalamic preoptic area MERFISH dataset^36^ (transcriptomic modality available), stGCL was able to achieve spatial vertical alignment of five consecutive tissue slices (Bregma-0.04, Bregma-0.09, Bregma-0.14, Bregma-0.19, and Bregma-0.24; **Fig. 4D**), identifying spatial domain boundaries highly similar to manual annotations (**Fig. 4E**). In contrast, GraphST and STAGATE failed to separate coherent tissue domains, and SCANPY could not decipher biologically meaningful structural domains other than the third ventricle (V3) region (**Fig. 4E and Supplementary Fig. S6**).

Stereo-seq is an emerging spatial transcriptomics platform that achieves subcellular spatial resolution and generates high-throughput ST datasets on a large number of cells. We evaluated the spatial clustering performance of stGCL using Stereo-seq data (transcriptomic modality available) from mouse olfactory bulb^9^ and mouse embryo samples^9^. The laminar structure of the coronal mouse olfactory bulb consists of the olfactory nerve layer (ONL), glomerular layer (GL), external plexiform layer (EPL), mitral cell layer (MCL), internal plexiform layer (IPL), granule cell layer (GCL), and rostral migratory stream (RMS) (**Fig. 4F**). stGCL accurately identified the outer layer distribution of organs (ONL, GL, EPL, and MCL) that matched with annotated layers. In the inner layer structures of the mouse olfactory bulb, it identified clusters (IPL, GCL, and RMS) with more accurate laminar distribution (**Fig. 4G, Supplementary Fig. S7A**). We further visualized marker genes in specific anatomical regions and discovered a strong correspondence between the clusters detected by stGCL and known marker genes, validating the clustering performance of stGCL (**Supplementary Fig. S7B**). In addition, we applied stGCL to four mouse embryos datasets acquired at the time stages of E9.5, E10.5, E11.5 and E12.5. stGCL provides intricate representations of tissue organization and organ distribution, closely resembling known anatomical annotations (**Supplementary Fig. S8**). Additionally, stGCL consistently and accurately identified the heart region across all four datasets. These results demonstrate that stGCL is capable of effectively deciphering Stereo-seq data.

Next, we tested the performance of stGCL on the Slide-seqV2 dataset of the mouse hippocampus, which contains gene expression profiles with approximate single-cell resolution^37^. We found that stGCL, GraphST and STAGATE are more effective than SCANPY in characterizing spatial domains and demonstrated clearer spatial separation (**Fig. 4H and Supplementary Fig. S9A**). It is noteworthy that stGCL is the only method capable of accurately identifying the “cord-arrow-like” structure in the hippocampal region. The “arrow-like” structure corresponds to the DG region, while the “cord-like” structure corresponds to the CA1, CA2, and CA3 regions of the Ammon’s horn. The division of CA1, CA2, CA3, and DG clusters in hippocampal structure by stGCL corresponds to the annotations in the Allen Reference Atlas and has been validated on several known marker genes^24,38^ (**Fig. 4H and Supplementary Fig. S9B**). For example, the secreted synaptic organizer molecule C1ql2 is a marker gene for dentate gyrus granule cells and is highly expressed in the identified DG domain^39^.

Further, we conducted tests on the mouse medial prefrontal cortex (mPFC) STARmap dataset, which features single-cell resolution. This dataset is composed of transcriptional and histological profiles and annotated as cortical layers L1, L2/3, L5, and L6^5^. Compared to the other methods, stGCL achieves the highest ARI, which means that the spatial domains identified by stGCL are mostly consistent with the original annotations (**Fig. 4I**). In addition, the domains detected by stGCL exhibit clear boundaries and less noise (**Fig. 4I and Supplementary Fig. S10A**).

Finally, we verified the generalization ability of stGCL on human pancreatic ductal adenocarcinoma (PDAC) ST data (spot resolution) and human breast cancer 10x Xenium data (subcellular resolution), both of which include gene expression and histological modalities. stGCL captures PDAC spatial domains that aligned better with the tissue structure provided by the original study (**Fig. 4G and Supplementary Fig. S10B**). Additionally, stGCL demonstrates spatial domains highly similar to the expected ductal carcinoma in situ (DCIS, named here DCIS #1 and #2), and invasive tumor regions in the breast cancer dataset (**Fig. 4K and Supplementary Fig. S10C**).

All these results demonstrate that stGCL is capable of analyzing spatial transcriptomic data generated on different sequencing platforms, even if these data have varying levels of spatial resolution and different available modalities. By effectively utilizing transcriptional profiles, histological profiles, and spatial information, stGCL enhances the accuracy and reliability of data interpretation, leading to the most precise and insightful biological outcomes.

### stGCL reveals intratumoral spatial heterogeneity from in-house bronchiolar adenoma dataset

Bronchiolar adenoma (BA) is a rare tumor that occurs in the bronchiolar epithelium of the peripheral lung and is known to potentially contain driver gene mutations found in lung cancer^40^. Currently, spatial transcriptomics studies on this type of tumor are relatively scarce. Therefore, we collected fresh tumor tissue to sequencing and engaged professional pathologists to manually segment the regions (see methods). The pathologist divided the BA tissue into tumor, normal, bronchus and blood vessel regions based on morphological features (**Fig. 5A**). The domains detected by stGCL are more consistent with manual annotation (ARI = 0.62) and have better regional continuity and less noise (**Fig. 5B**). For example, stGCL and GraphST display clear spatial patterns, whereas GraphST is unable to identify the tiny tumor region at the top part. The results from STAGATE, SpaGCN and SCANPY show difference from manual annotation, with fragmented irregular boundaries between domains. We applied PAGA algorithm to infer associations between tissue domains and the embedding from stGCL shows closer connections among normal lung tissue, blood vessel and bronchus (**Fig. 5C**).

**Fig. 5.**
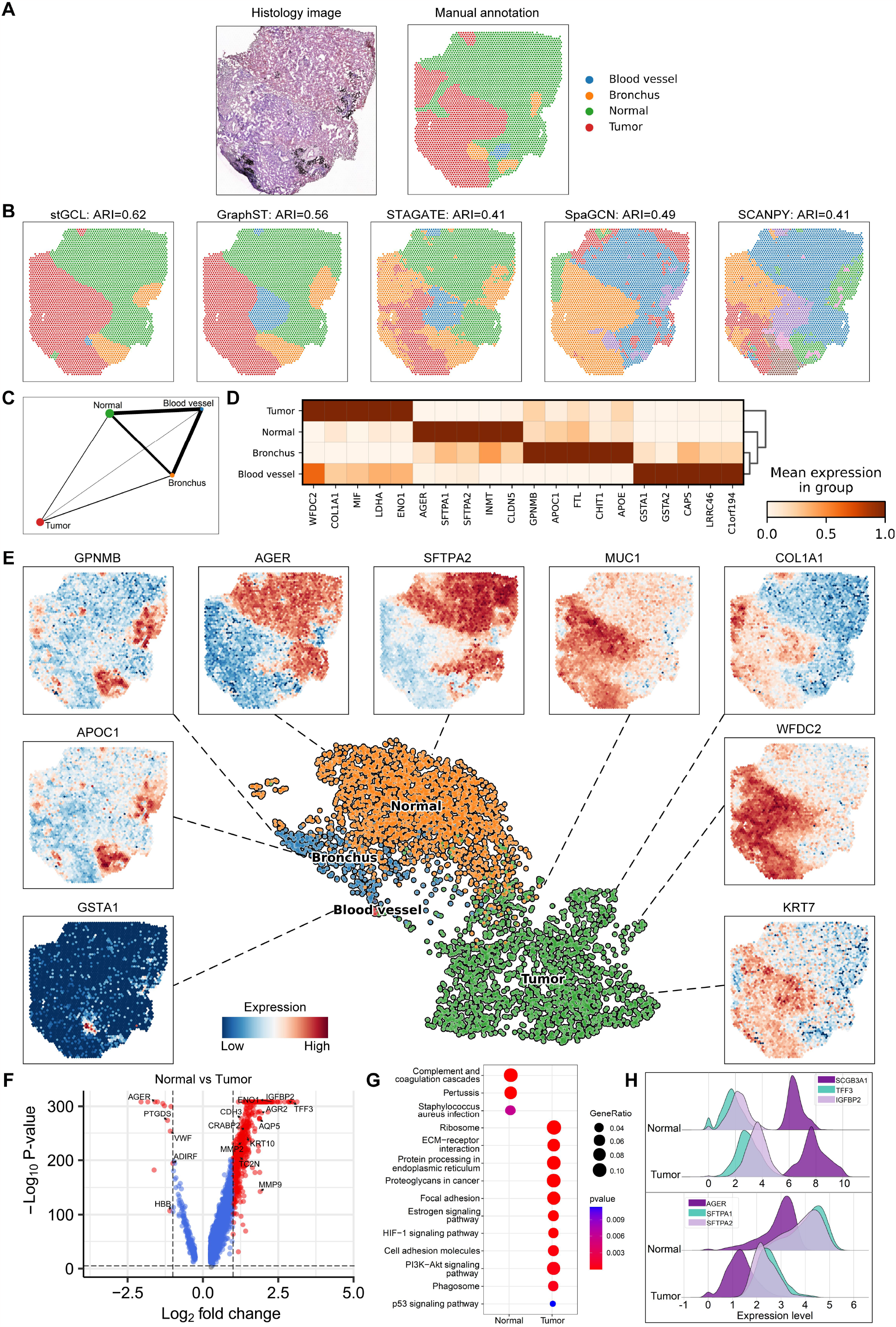
stGCL effectively distinguishes intratumoral spatial heterogeneity in bronchiolar adenoma. **A**. H&E stained image and manual annotation of tissue slice by a pathologist. **B**. Clustering results of stGCL, GraphST, STAGATE, SpaGCN and SCANPY. **C**. Trajectory inference results using PAGA on stGCL embeddings. **D**. Heatmap of the top 5 DEGs for the 4 domains of stGCL. Rows and columns denote domains and genes, respectively. **E**. UMAP visualization and scatter plot (Tumor: MUC1, KRT7, COL1A1 and WFDC2; Normal: AGER and SFTPA2; Bronchus: GPNMB and APOC1; Blood vessel: GSTA1) generated by stGCL. **F**. Volcano plot of DEGs between tumor and normal regions. **G**. KEGG pathways of DEGs between tumor and normal regions. **H**. Ridge plots of log-normalized gene expression of top DEGs in tumor and normal regions.

To better understand the relationship between the distribution of different tissue regions and their biological functions, we conducted a detailed analysis of differentially expressed genes (DEGs) in the domains identified by stGCL. Initially, we created the Heatmap and scatter plot for the differentially expressed genes within each region. The results demonstrated that DEGs are well segregated in different tissue regions, serving as domain-specific marker genes (**Fig. 5D, E**). In the tumor region, MUC1, COL1A1, WFDC2 and KRT7 genes exhibited high expression (**Fig. 5E**). MUC1 and KRT7 are reported to be predefined BA markers, and they are positive in BA^41,42^. COL1A1 is a marker of fibroblasts and a key gene in the development and progression of lung cancer, which is associated with tumor cell migration, invasion and proliferation^43,44^. The expression level of WFDC2 is significantly increased in lung cancer tissues, and WFDC2 is a potential biomarker for early diagnosis of lung cancer^45^. In addition to the aforementioned results that align with previous studies, we also made new discoveries. We found that COL1A1 and WFDC2 exhibited significant expression differences between normal and tumor tissues, suggesting that they may play a crucial regulatory role in the occurrence and development of BA. Additionally, the results indicate that AGER is highly expressed in human lung tissue, particularly in alveolar epithelial cells, and it possesses a latent function in epithelial-extracellular matrix interactions^46^. Similar to AGER, SFTPA2 exhibits persistently high expression in the lung, and together with SFTPA1-encoded human surfactant protein A (SP-A), they play a critical role in lung homeostasis and immunity^47^. For the bronchus region, GPNMB encodes a type I transmembrane glycoprotein and it is expressed in bronchial epithelial cells^48^.

To further unveil the gene expression and functional differences between normal lung tissue regions and tumor tissue regions, we conducted a more detailed differential expression analysis and pathway enrichment analysis for these two areas (**Fig. 5F-H**). Compared with normal lung tissue regions, some of the highly expressed genes in tumor regions were associated with lung cancer risk (**Fig 5F, H**). For example, the matrix metalloproteinase (MMP) gene family members MMP2 and MMP9 and mesenchymal marker genes play a major role in tumor invasion through proteolysis of the extracellular matrix. They also promote tumor growth and angiogenesis^49^. SCGB3A1 is related to cell differentiation and proliferation as a lung cancer promoter^50^, and IGFBP2 is a pleiotropic oncogenic protein highly expressed in tumor tissues^51^. In addition, TFF3 is associated with tumor metastasis and prognosis and can be used as a novel biomarker for lung cancer^52^. We further found that these DEGs are significantly involved PI3K-Akt signaling pathway, ECM-receptor interaction, Proteoglycans in cancer (**Fig 5G**). Most of the DEGs and pathways found in tumor regions are associated with lung cancer, which fits well with the biological insight that BA may contain driver gene mutations found in lung cancer.

The above analysis results demonstrate that using stGCL to analyze spatial transcriptomics data of BA can reveal gene expression differences in different regions of BA tissue, providing a deeper understanding of the spatial heterogeneity of the tumor. Additionally, stGCL has identified new potential biomarker target genes, such as COL1A1 and WFDC2. This is significant for exploring the origins and development of BA, identifying key biomarkers and therapeutic targets, and advancing personalized medicine.

## Discussion

Delving deeply into the rich multi-modal information within ST data is crucial for understanding the organization, function and disease progression of heterogeneous tissues. In this work, we proposed stGCL, a cross-modality fusion model based on multi-modal graph contrastive learning designed to incorporate multi-modal data and generate precise low-dimensional embeddings, effectively enabling spatial domain detection, multi-slice integration, and related downstream analyses. stGCL introduces a novel H-ViT method to learn the histological features of each spot, followed by the adaptive aggregation of transcriptional and histological profiles with spatial neighborhood information via multi-modal GATE. It then incorporates global information from tissue slices as a guidance signal and utilizes contrastive learning to further enhance joint embeddings of spots. Compared to other methods, stGCL demonstrates superior accuracy in analyzing spatial transcriptomic data from different platforms and has the capability to integrate multiple slices.

The prominent advantage of stGCL in the field of ST data analysis is manifested in the multifunctionality of its generated latent embeddings. These embeddings are capable of not only accurately identifying spatial domains but also integrating with other advanced bioinformatics algorithms to conduct a range of complex and in-depth analyses. This includes differential expression analysis, intercellular communication, and developmental trajectory inference. For instance, in the analysis of the DLPFC dataset, the combination of stGCL with the PAGA algorithm precisely identified the developmental trajectory from Layer 1 to Layer 6. In the analysis of the bronchiolar adenoma dataset, stGCL identified novel potential biomarkers, offering possibilities for early diagnosis and treatment of the disease. This multifunctionality enables stGCL to deeply dissect the gene expression differences and spatial heterogeneity of heterogeneous tissues, while also revealing richer and more profound biological information. The flexibility and multifaceted analytical capabilities of stGCL significantly broaden its application in biomedical research. It provides robust support for understanding disease progression, identifying key biomarkers, discovering therapeutic targets, and advancing personalized medicine.

Although stGCL has demonstrated exceptional performance in its current stage, there remains room for improvement. The H-ViT model that stGCL relies on is pre-trained on the ImageNet dataset and has not been specifically fine-tuned for the characteristics of histology images. This may limit its capability in mining spot histological features.

With the rapid advancement of spatial transcriptomics technology, there is a significant increase in spatial resolution and data throughput, necessitating computational methods for ST data to meet higher memory efficiency and time efficiency requirements. To address this, we conducted tests on the runtime and GPU memory consumption of stGCL on both real and simulated datasets (**Supplementary Fig. S11**). The test results indicated a direct proportional relationship between the runtime and GPU memory usage with the number of spots. Additionally, incorporating histology image analysis was observed to further increase the demands on runtime and GPU memory. Notably, on a breast cancer dataset, generated using the advanced 10x Xenium technology and containing over 167k cells, stGCL only required approximately 14 minutes for processing and consumed about 26.9GB of GPU memory on a server with Intel(R) Xeon(R) Gold 6258R CPU @ 2.70GHz and NVIDIA QuADro GV100 GPU (**Supplementary Fig. S11A**). This result clearly demonstrates the capability of stGCL to handle large-scale ST data. In light of these findings, we believe that stGCL will prove to be a competitive analysis method in the future domain of subcellular resolution ST data analysis.

In summary, stGCL is a robust and widely applicable multi-modal data fusion algorithm for ST data. It is capable of efficiently processing single or multiple biological tissue slices, as well as effectively integrating and analyzing multi-modal data. stGCL is poised to become a powerful tool in the field of ST research, offering substantial support in understanding disease progression and identifying key biomarkers. This algorithm not only facilitates advanced analytical capabilities but also promises to significantly contribute to the advancements in the study of complex biological systems and diseases.

## Methods

### ST sequencing data preprocessing

We employed the SCANPY package for processing the raw gene expression counts^21^. This process included log-transformation, normalization according to library size, and scaling to achieve unit variance and zero mean. Subsequently, the top 3000 highly variable genes (HVGs) were selected. These HVGs were then used to form the initialized gene expression matrix, which was inputted into stGCL.

### Encoding histology image features using H-ViT

The texture (cell size, shape, and arrangement) and color of histology images profoundly reveal the heterogeneity of biological tissues. For ST data with histology images, we proposed a modified ViT^53^ model, H-ViT, which is designed to effectively extract histological features and remove noise in tissue slices (**Fig. 1B**). Specifically, given a histology image **I**_*h*_, it can be split into *n* spot images based on the size and position of each spot. Next, the image **I**_*i*_ ∈ ℝ^*h*×*w*×*c*^ corresponding to each spot *i* is divided into image patches, where (*h, w*) denotes the resolution of the spot image and *c* = 3 denotes the number of channels. We then flatten all patches and assemble them into a *N* ×(*p*^2^ ×3) sequence **I**_*p*_, where (*p, p*) represents the resolution of each image patch and *N* = *hw p*^2^ represents the number of patches. In this study, we set *h* = *w* = 40 pixels. For each spot image **I**_*i*_, the main steps of the model are as follows:

1. Patch embedding: To extract the features of each spot image, we employ a simple fully connected layer **W**_*p*_ to map the sequence matrix **I**_*p*_ to the patch embedding **E**_*p*_ ∈ ℝ^*N*×768^, i.e., **E**_*p*_ = **I**_*p*_ · **W**_*p*_. The obtained embedding matrix is used as the input of the Transformer encoder.
2. Transformer encoder: The core operation of the Transformer is multi-head self-attention (MSA), which can adaptively learn the attention of patches sequence in the spot image. MSA is a linear combination of attention heads:

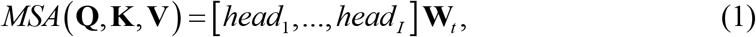

where **W**_*t*_ denotes a learnable weight matrix for aggregating attention heads. **Q**, **K** and **V** are query, key and value. The attention mechanism can be expressed as:

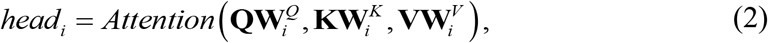

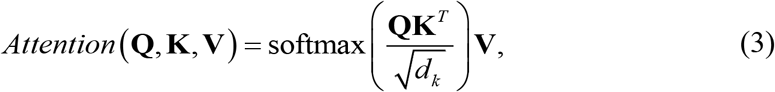

where 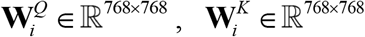 and 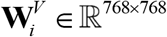 refer to learnable weight matrices. In the equation (3), the first term 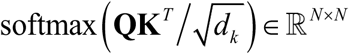 is called the Attention Map, each column of it indicates the attention weights contributed by other elements in the sequence. The second term **V** denotes the value of the self-attention mechanism. The Transformer encoder is composed of MSA and MLP blocks, and the output is an *N* × 768 matrix. In the end, H-ViT encodes the histology image into a *n* ×(*N* ×1000) matrix as the final output. To better capture the histological information and balance it with gene expression information, principal component analysis (PCA) was then employed to extract the first 3000 principal components (PCs) on the output matrix to obtain the histological feature coding matrix **I**. In each ST data, number of attention heads and MSA layers of the H-ViT model are 12 and 6, respectively.

### Building the spatial neighborhood graph

In ST data, spatial coordinate information can reflect the potential relationship between spots. Therefore, we model the spatial information as an undirected graph *G* = (*V, E*) based on a pre-defined radius *r*. In graph *G*, a node corresponds to a spot, and an edge connects a pair of spots. We use the image coordinates to compute the Euclidean distance between spots, which in turn builds an adjacency matrix **A** ∈ ℝ^*n*×*n*^. **A**_*ij*_ = 1 if the Euclidean distance between spot *i* and spot *j* is less than *r*, otherwise 0. In experiments, we set *r* ∈ (*d*, 2*d*) (*d* is the spacing between adjacent spots), which achieves the best performance on most ST data. The self-loop is added to each spot.

### Multi-modal graph contrastive learning for representation enhancement

The proposed stGCL is a cross-modal fusion framework that uses multi-modal graph contrastive learning as its core algorithm, which takes preprocessed gene expression, encoded histological features and constructed spatial neighbor graph as inputs to learn latent spot embeddings (**Fig. 1A)**. The stGCL is structured into three primary phases: 1) Data Augmentation; 2) Multi-Modal Graph Attention Auto-Encoder for Structured Embedding; and 3) Contrastive Learning for Enhanced Spot Representations. Detailed descriptions of the implementation for each of these steps are provided subsequently.

#### Data Augmentation

Data augmentation, a pivotal technique to amplify sample size and mitigate model overfitting, plays a vital role in graph contrastive learning. In the context of spatial transcriptomics data, this approach lends itself well to the natural modeling of ST data as graph-structured data. Within this framework, each spot node is characterized by two distinct attributes: gene expression and histological features, the latter being included when available. This dual-attribute configuration forms the foundation of our data representation in the proposed framework. We use the corruption function *C* to generate a negative sample, 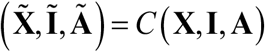. In particular, *C* randomly shuffles the rows of the gene expression matrix and histological feature matrix, and keeps the spatial neighbor graph structure unchanged.

#### Multi-Modal Graph Attention Auto-Encoder for Structured Embedding

In this work, we present an extension to the GATE^54^ model, leading to the development of the multi-modal GATE. stGCL employs this multi-modal GATE to effectively learn joint latent features. This is achieved by integrating data from different modalities, specifically histological features (when available) and gene expression data. This integration process facilitates a comprehensive understanding and representation of the underlying biological processes, leveraging the strengths of each modality to enrich the learned feature space. Specifically, multi-modal GATE adopts graph attention networks (GAT)^55^ as the encoder to learn the spot joint structured embedding **H** by iteratively aggregating the features of neighbor nodes. The inputs to the encoder are the gene expression matrix **X**, the histological feature matrix **I**, and the adjacency matrix **A**, For each modality, the *l* -th encoder layer learns the latent representation of the spot *i* (*i* ∈{1, 2,…, *n*}) as follows:

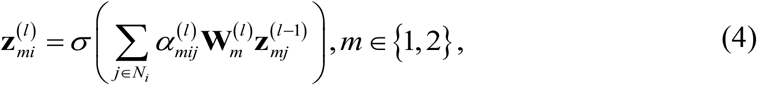

where 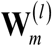 and *σ* represent the learnable weight matrix and nonlinear activation function, respectively. *m* is the number of modalities, and *N*_*i*_ is the neighbors of the spot *i* in the graph *G*. 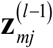 is the representation of the spot *j* generated by the *l* −1 -th encoder layer in the *m* -th modality. 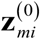 is initialized as gene expression profile 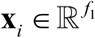 or histological profile 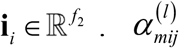 denotes the normalized attention coefficient output by the *l* -th graph attention layer in the *m* -th modality. We then initialize the joint embedding 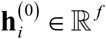 by performing a concatenation operation on the latent representations 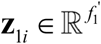 and 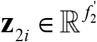 of each modality, where 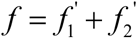. The joint structured embedding of spot *i* is defined as

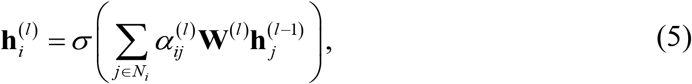

where 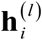 is the embedding of the *l* -th layer. The final output 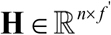 of the encoder is the spot joint embedding, which simultaneously learns gene expression information, histological information and spatial information. 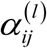 represents the normalized attention coefficient output by the *l* -th graph attention layer. The graph attention layer incorporates a self-attention mechanism to dynamically evaluate the relevance between individual spots and their neighboring nodes. Each graph attention layer is computed by

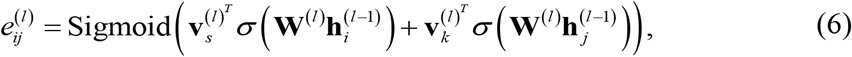

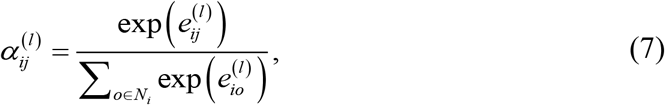

where 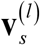 and 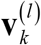 denote the trainable parameters. Sigmoid is the sigmoid function. 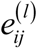 represents the edge weight between the spot *i* and its neighbor spot *j* of the *l* -th graph attention layer. Similarly, we can input 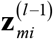 and 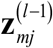 to calculate 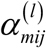 in the graph attention layer. This attention-driven mechanism ensures a more nuanced and context-aware representation of the spatial data, enhancing the overall learning process.

The decoder part of the multi-modal GATE takes the spot joint embedding **H** as input, and each decoder layer reverses the process of its corresponding encoder layer. In this work, the decoder and encoder form a symmetric architecture (**Fig. 1A**). The spot features of the *l* −1-th layer are reconstructed by the *l* -th layer decoder:

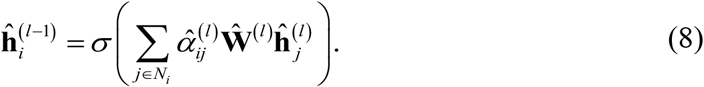

After that, we split the output 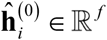 into 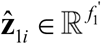 and 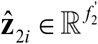 as the input to the decoder for each modality. Features of gene expression profile and histological profile are reconstructed by the decoder

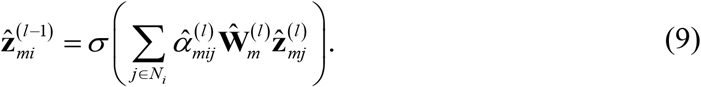

stGCL employs half of the trainable parameters (i.e. 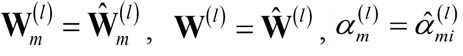 and 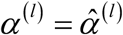) to avoid overfitting. For gene expression profile and histological profile, we train multi-modal GATE by minimizing the reconstruction loss of spot features:

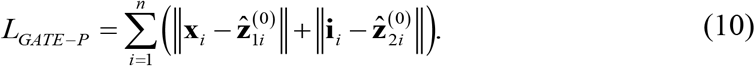

For the negative sample, we also use the above multi-modal GATE model to accurately learn the spot joint low-dimensional representation 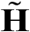. The overall reconstruction loss for multi-modal GATE is summarized as

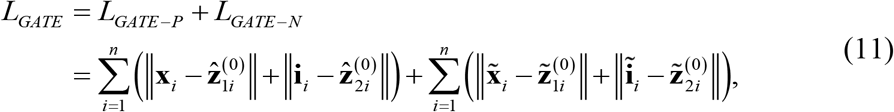

where 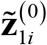 and 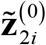 represent modality-specific features obtained by dividing 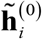 into two latent vectors.

#### Contrastive Learning for Enhanced Spot Representations

In spatial transcriptomics data derived from the same tissue slice, the gene expression and histological features of individual spots generally align with the global features of the entire tissue slice^56^. This inherent consistency provides a supervised signal that we utilize to guide the learning of spot joint embeddings, thereby enhancing their quality and discriminative power. Specifically, stGCL is designed to maximize the mutual information between each spot’s joint embedding and the global summary of the graph. This strategic approach ensures that the learned joint representations capture both local attributes, such as spot-specific expression patterns, and global characteristics, like the overall tissue microenvironment patterns. We employed a readout function 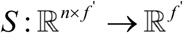 to yield the graph-level summary **s** = *S* (**H**) by following Velickovic et al^57^. The negative samples 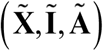 obtained by data augmentation are then fed into the multi-modal GATE to produce the spot joint representation 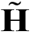. Furthermore, we evaluate the spot embedding-summary pair using the discriminator 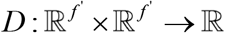, and a higher probability score indicates that the spot embedding is more likely to be included in the summary. At the same time, the probability score of positive pair (**h**_*i*_, **s**) is higher than that of negative pair 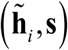. Finally, the objective function of contrastive learning adopts binary cross-entropy (BCE), which is defined as follows

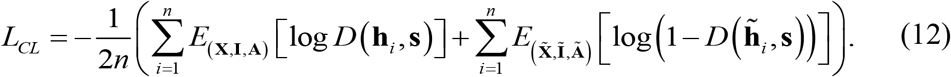

The overall loss function of stGCL is a combination of multi-modal GATE reconstruction loss and contrastive loss like below:

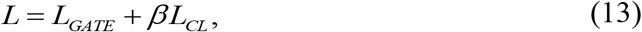

where *β* is the trade-off parameter to control the influence of the contrastive loss term, and its default value is 0.04. In stGCL, the encoder and decoder of the multi-modal GATE module are both a two-layer GAT. We set 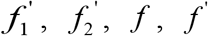 to 100, 100, 200, 30 respectively, and choose exponential linear unit (ELU)^58^ as the activation function. The Adam algorithm^59^ is adopted to optimize the objective function *L* of stGCL. The learning rate, weight decay and training epoch are set to 1e-4, 1e-4 and 1200 (default), respectively. Implementation details are presented in the supplementary material. For more details, please see **Supplementary Fig. S12**.

### Spatial clustering and visualization

Leveraging the stGCL embeddings **H**, we employ either the mclust^60^ algorithm or the Louvain algorithm^10^ to cluster the spots into distinct spatial domains. Each identified cluster represents a spatial domain, encompassing spots that share not only similar expression and histological profiles but also exhibit spatial proximity. In scenarios where the number of spatial domains is predefined, the mclust algorithm is utilized, setting the number of clusters equal to the number of actual labels. In the absence of prior information regarding the ST data, we opt for the Louvain algorithm.

After clustering, it is acknowledged that certain spots may be erroneously classified into different spatial domains. To address this, stGCL introduces an optional refinement step. Specifically, for a given spot, stGCL initially examines the spatial clustering outcomes of the spot and its proximal neighbors (typically the 50 nearest neighbors). Following this, stGCL reassigns the spot to the cluster that corresponds to the most frequent label among these neighbors. It is important to note that this refinement step is applied exclusively to datasets where the identification of spatial domains is pertinent and is not recommended for datasets aimed at cell type detection.

For ST datasets with manual annotations, we utilize the Adjusted Rank Index (ARI)^61^ to assess the clustering efficacy of each method, with higher ARI values indicating superior performance. Additionally, the Kuhn-Munkres^62^ algorithm is employed to align cluster labels with ground truth labels effectively. For visualization purposes, the UMAP method is used to project the joint embedding matrix **H** into a two-dimensional space, facilitating easier interpretation and analysis of the clustering results.

### Integration of multiple tissue slices via 3D spatial structure reconstruction

The integration of multiple tissue slices offers invaluable insights into the overall structural organization and functional dynamics of the tissue. To capitalize on this, we have enhanced the stGCL method to support the joint analysis of multiple tissue slices. This extension enables a more thorough exploitation of the shared information and spatial context across different slices. Our proposed alignment algorithm is designed to align spatial coordinates from adjacent horizontal slices and consecutive vertical slices, as depicted in **Fig. 1C**. By aligning these spatial coordinates, we are able to construct a spatial neighborhood graph for multiple slices in a manner analogous to that used for a single slice. This comprehensive graph not only incorporates neighbor information within and across multiple slices but also effectively smoothens spot features on the cut surfaces, thus mitigating batch effects. The gene expression data, histological features, and the spatial neighborhood graph derived from multiple slices are then input into the stGCL framework, following the same processing steps as outlined for single-slice stGCL analysis. The resultant joint embeddings from this analysis of multiple slices are then utilized to identify spatial domains. This process significantly enriches our understanding of tissue organization and functionality, providing a more holistic view than what could be achieved by analyzing single slices in isolation.

To reconstruct the original 3D spatial structure of the tissue, we executed both vertical and horizontal alignments of the slices. Operating under the assumption that the cut surfaces of the slices are nearly identical in shape and size, we developed an alignment algorithm focusing on the edge structure of these surfaces. This method facilitates the integration of multiple slices by aligning their edge structures as closely as possible. For consecutive slices, the cut surface encompasses the entire slice, providing a comprehensive framework for alignment. In the process of vertical integration, we designate a reference tissue slice (referred to as section 1) and then modify the coordinates of the remaining slices to align with this reference. Assuming there are *s* slices, we calculate the mean of the spot coordinates (*x*_*i*_, *y*_*i*_) located on the edge structures of all slice cut surfaces, thus determining the center spot coordinate 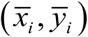 (*i* ∈{1, 2,…, *s*}). Then, we compute the bias between the central spot coordinates of section 1 and those of the other sections 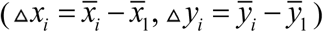. This bias calculation enables us to adjust the positions of the other sections, achieving two-dimensional (2D) alignment across multiple tissue slices. Following the 2D alignment, we proceed to establish the z-axis coordinates for all tissue slices, thereby extending the spatial representation from 2D within each slice to a 3D space encompassing multiple slices. This step is crucial for accurately reconstructing the tissue’s original 3D spatial structure and understanding its organization. In particular, the z-axis coordinates of the spots in the -th slice can be defined as

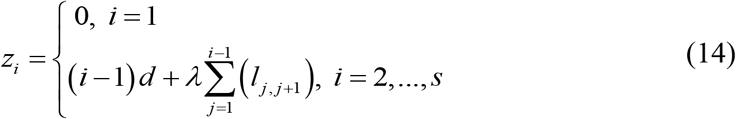

where *d* is the thickness of the tissue slice and *l*_*j, j*+1_ is the separation distance between slice *j* and slice *j* + 1. For instance, the 10x Visium platform generates tissue slices that have a thickness ranging from 10 to 20 μm, which is notably thinner than the diameter of each spot, measured at 55 μm^19^. Given this disparity, it becomes imperative to carefully consider the neighboring spots between consecutive slices, which are constructed based on aligned spatial coordinates and a predefined radius, as illustrated in **Fig. 1C**.

In the case of horizontal integration, we specifically focused on translating and aligning adjacent tissue slices along the x- and y-axis directions. In this context, the cutting line within a tissue slice is designated as the y-axis, while its perpendicular counterpart is defined as the x-axis. For any given pair of slices, we identify the rightmost spots in the left slice (referred to as section 1) and the leftmost spots in the right slice (section 2), considering these as spots situated on the edge structure of the cutting surface, as depicted by the yellow spots in **Fig. 1C**.

For horizontal integration, we translated and aligned adjacent tissue slices from the x- and y-axis directions, respectively. The cutting line in a tissue slice is called the y-axis, and perpendicular to it is the x-axis. For a given two slices, we consider the rightmost spots in the left slice (section 1) and the leftmost spots in the right slice (section 2) as spots on the edge structure of the cutting surface (yellow spots in **Fig. 1C**). Our initial step involves calculating the mean value 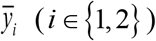 of the y-axis coordinates of the spots on the edge structures of both slices. We then determine the maximum value (max (*x*_1_)) of the x-axis coordinates of all spots in section 1 and the minimum value (min (*x*_2_)) of the x-axis coordinates of all spots in section 2. Finally, we fix the position of section 1 and adjust the coordinates of section 2 based on the calculated bias 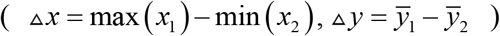 to achieve horizontal integration. Furthermore, this approach allows us to treat the aligned slices as a single, unified entity. We can then apply the aforementioned horizontal integration method to align additional slices in various directions.

### Spatial trajectory inference

Trajectory inference represents a significant methodology for deducing the dynamic developmental processes of cells. Utilizing the joint embedding derived through stGCL, we employ the PAGA algorithm^22^, sourced from the SCANPY package (‘scanpy.pl.paga_compare’ function)^21^, to elucidate the spatial trajectory of these cellular processes.

### Identifying differentially expressed genes

Differentially expressed genes (DEGs) exhibit unique expression patterns across various spatial domains. To identify these DEGs, we perform differential expression analysis utilizing the Wilcoxon test implemented within the SCANPY package. Genes that meet a stringent criterion of a 1% false discovery rate threshold are selected as DEGs, ensuring high confidence in the differential expression results.

### Cell-cell communication

Cell-cell communication is fundamental to cellular function and tissue homeostasis. Leveraging the cluster predictions derived from stGCL embeddings, we utilize the CellChat^33^ method to infer ligand-receptor (L-R) interactions among cells.

### Sample Collection

Fresh tumor tissue was obtained from a patient diagnosed with primary bronchiolar adenoma (BA), who underwent surgical resection at the Shandong Cancer Hospital and Institute. Prior to surgery, the patient had not received any form of neoadjuvant therapy.

### Ethics declarations

This study received approval from the Ethics Committee of Shandong Cancer Hospital and Institute, under the reference number SDTHEC2021012059. Informed consent was obtained from the donors.

### Spatial transcriptome sequencing

This experiment utilizes the Visium Technology Platform developed by 10x Genomics company. The reagents and consumables used in the study are supplied by this platform, and you can find the corresponding product numbers by visiting https://www.10xgenomics.com/products/spatial-gene-expression.

### Manual region segmentation

For the bronchiolar adenoma sample, the pathologist used labelme software^63^ to annotate the histology image into distinct regions, each of which can be considered as an irregular polygon. We designed a strategy to identify which region each spot is located in. With spot *i* as the endpoint, we can make the ray *l*_*I*_ in any direction. Then we employ the OpenCV package^64^ to determine whether the ray passes through the polygon. Which region the spot belongs to can be determined based on the number of times the ray crosses the polygon boundary (crossing number). We can conclude that the spot *i* is in a polygon *p* if the crossing number between the ray and polygon *p* is odd. On the contrary, the spot *i* is outside polygon *p* if the crossing number between the ray and polygon *p* is even. The formula is defined as follows:

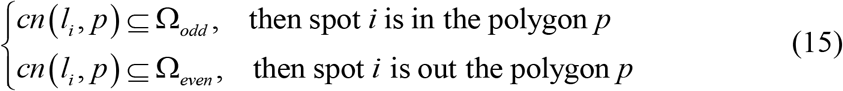

where *cn* (*l*_*i*_, *p*) is the crossing number between the ray *l*_*I*_ and polygon *p*, and Ω_*odd*_ means odd number set, Ω_*even*_ means even number set. The proposed manual region segmentation strategy greatly facilitates researchers in automatically converting the annotated information provided by pathologists into label information for each spot. The manual region segmentation strategy we propose significantly aids researchers in the automated conversion of pathologist-provided annotated information into specific label data for each spot. This strategy is pivotal in enabling the transformation of unlabeled datasets into accurately labeled ST data.

### Dataset description

In this study, we utilize 11 ST datasets sourced from diverse platforms, including 10x Visium, STARmap, MERFISH, Slide-seqV2, NanoString CosMx SMI, ST, 10x Xenium, and Stereo-seq, to validate the efficacy of our proposed method. This selection of datasets represents a broad spectrum of technological approaches in the field of spatial transcriptomics, thereby providing a comprehensive basis for assessing the performance and versatility of our method across various platforms.

The dorsolateral prefrontal cortex (DLPFC) 10x Visium dataset is derived from 12 tissue slices of three human brains, each slice comprising 3,460 to 4,789 spots and is manually annotated as DLPFC layers and white matter (WM)^20^.

Mouse brain dataset profiled by 10x Visium is divided into anterior and posterior sections. The mouse anterior brain dataset contains 2,695 spots and 32,285 genes, and is manually partitioned into 52 regions^15^.

The mouse olfactory bulb Stereo-seq dataset with sub-cellular level resolution contains 19,109 bins and 14,376 genes^9^.

The mouse embryo dataset from the Stereo-seq platform consists of slices (E9.5: 5,913 bins; E10.5: 18,408 bins; E11.5: 30,124 bins; E12.5: 51,365 bins) from four developmental stages^9^.

The mouse medial prefrontal cortex (mPFC) STARmap dataset with single-cell resolution maps 1,053 cells with 166 genes and is divided into the four distinct layer structures by the original study^5^.

The mouse hypothalamus MERFISH dataset is obtained from tissue slices at Bregma-0.04, -0.09, -0.14, -0.19 and -0.24 mm in the preoptic area of the hypothalamus of animal 1^36^. Five tissue slices contain 5,488, 5,557, 5,926, 5,803, and 5,543 cells, respectively, and are segmented into eight brain structures including the third ventricle (V3), bed nuclei of the strata terminalis (BST), columns of the fornix (fx), medial preoptic area (MPA), medial preoptic nucleus (MPN), periventricular hypothalamic nucleus (PV), paraventricular hypothalamic nucleus (PVH), and paraventricular nucleus of the thalamus (PVT)^65^.

The approximate single-cell resolution mouse hippocampus data obtained from the Slide-seqV2 platform consists of 52,869 spots^37^.

The human non-small cell lung cancer (NSCLC) dataset with subcellular level resolution was captured by NanoString CosMx SMI technology^32^. It includes 20 FOVs with a total of 82,843 cells and 12 cell types.

The human pancreatic ductal adenocarcinoma (PDAC) dataset obtained from the ST platform measured 19,738 genes in 428 spots and is manually pathologically annotated into 4 regions^66^.

The human breast cancer 10x Xenium dataset contains 167,780 cells with 3 manually annotated tissue regions^67^.

The in-house bronchiolar adenoma (BA) dataset profiled by 10x Visium contains 4,002 spots and 36,601 genes, and is manually partitioned into 4 regions. These datasets are summarized in Supplementary Table S1.

## Data availability

The public datasets used in this paper are freely available. (1) Human dorsolateral prefrontal cortex (DLPFC) 10x Visium dataset (http://spatial.libd.org/spatialLIBD/); (2) Mouse brain section 1 10x Visium dataset (https://www.10xgenomics.com/resources/datasets/mouse-brain-serial-section-1-sagittal-anterior-1-standard-1-1-0, https://www.10xgenomics.com/resources/datasets/mouse-brain-serial-section-1-sagittal-posterior-1-standard-1-1-0) and mouse brain section 2 10x Visium dataset (https://www.10xgenomics.com/resources/datasets/mouse-brain-serial-section-2-sagittal-anterior-1-standard-1-1-0, https://www.10xgenomics.com/resources/datasets/mouse-brain-serial-section-2-sagittal-posterior-1-standard-1-1-0); (3) Mouse olfactory bulb Stereo-seq dataset (https://github.com/JinmiaoChenLab/SEDR_analysiss); (4) Mouse embryo Stereo-seq dataset (https://db.cngb.org/stomics/mosta/); (5) Mouse medial prefrontal cortex (mPFC) STARmap dataset (https://github.com/zhengli09/BASS-Analysis); (6) Mouse hypothalamus MERFISH dataset (https://datadryad.org/stash/dataset/doi:10.5061/dryad.8t8s248); (7) Mouse hippocampus Slide-seqV2 dataset (https://singlecell.broadinstitute.org/single_cell/study/SCP354/slide-seq-study); (8) Human non-small cell lung cancer (NSCLC) dataset (https://nanostring.com/resources/smi-ffpe-dataset-lung13-data/); (9) Human pancreatic ductal adenocarcinoma (PDAC) ST dataset (https://www.ncbi.nlm.nih.gov/geo/query/acc.cgi?acc=GSM3036911); (10) Human breast cancer 10x Xenium dataset (https://www.10xgenomics.com/products/xenium-in-situ/preview-dataset-human-breast).

The in-house dataset used in this study has been deposited on Zenodo, which is publicly accessible. (11) Human bronchiolar adenoma (BA) 10x Visium dataset (https://zenodo.org/record/8185216). Further information and requests for resources and raw FASTQ data should be directed and will be fulfilled by the Contact: R.G.

## Code availability

The open-source Python implementation of the stGCL toolkit is available on both GitHub (https://github.com/RuiGaolab/stGCL) and Zenodo (https://zenodo.org/record/8185216). Detailed manuals and tutorials are provided.

## Acknowledgements

This work was supported by the National Natural Science Foundation of China [No. U1806202 to R.G., No. 61973190 to Z.L., No. 62303271 to W.Z.], the National Key Research and Development Program of China [No. 2020YFA0712402 to Z.L.], the Program of Qilu Young Scholars of Shandong University, and the Fundamental Research Funds for the Central Universities [No. 2022JC008 to Z.L.], the Key Research and Development Program of Shandong Province [No. 2021CXGC010506 to W.L.]. The research was also supported with a project grant from the Crafoord Foundation (Crafoordska stiftelsen) grant to Y.D.M..

## Author contributions

R.G., Y.D.M. and W.L. conceived and supervised the project. N.Y. and D.Z. designed the model. D.Z. developed the stGCL software. N.Y., D.Z., R.G., W.Z. and Y.D.M. wrote the manuscript. N.Y., D.Z., R.G. led the data analysis with input from Z.L., X.Q., W.Z., and C.W. B.C. and W.L. provided the bronchiolar adenoma tissue, and M.Z. annotated this data. All authors read and approved the final manuscript.

