## Supplementary Figures and Tables for "stGCL: A versatile cross-modality fusion method based on multi-modal graph contrastive learning for spatial transcriptomics"

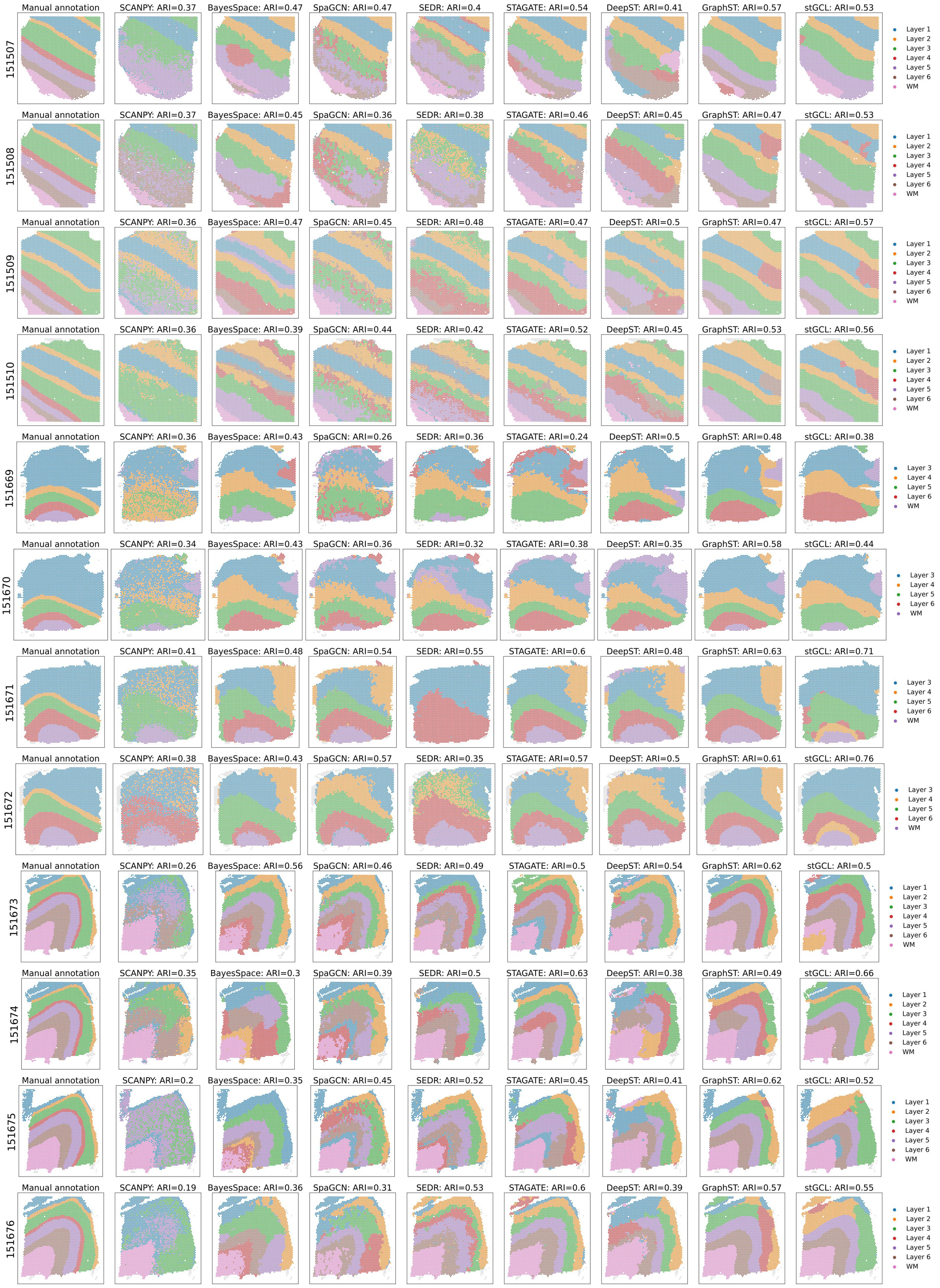
151507
Manual annotation
SCANPY: ARI=0.37
BayesSpace: ARI=0.47
SpaGCN: ARI=0.47
SEDR: ARI=0.4
STAGATE: ARI=0.54
DeepST: ARI=0.41
GraphST: ARI=0.57
stGCL: ARI=0.53
Layer 1
Layer 2
Layer 3
Layer 4
Layer 5
Layer 6
WM
151508
Manual annotation
SCANPY: ARI=0.37
BayesSpace: ARI=0.45
SpaGCN: ARI=0.36
SEDR: ARI=0.38
STAGATE: ARI=0.46
DeepST: ARI=0.45
GraphST: ARI=0.47
stGCL: ARI=0.53
Layer 1
Layer 2
Layer 3
Layer 4
Layer 5
Layer 6
WM
151509
Manual annotation
SCANPY: ARI=0.36
BayesSpace: ARI=0.47
SpaGCN: ARI=0.45
SEDR: ARI=0.48
STAGATE: ARI=0.47
DeepST: ARI=0.5
GraphST: ARI=0.47
stGCL: ARI=0.57
Layer 1
Layer 2
Layer 3
Layer 4
Layer 5
Layer 6
WM
151510
Manual annotation
SCANPY: ARI=0.36
BayesSpace: ARI=0.39
SpaGCN: ARI=0.44
SEDR: ARI=0.42
STAGATE: ARI=0.52
DeepST: ARI=0.45
GraphST: ARI=0.53
stGCL: ARI=0.56
Layer 1
Layer 2
Layer 3
Layer 4
Layer 5
Layer 6
WM
151669
Manual annotation
SCANPY: ARI=0.36
BayesSpace: ARI=0.43
SpaGCN: ARI=0.26
SEDR: ARI=0.36
STAGATE: ARI=0.24
DeepST: ARI=0.5
GraphST: ARI=0.48
stGCL: ARI=0.38
Layer 3
Layer 4
Layer 5
Layer 6
WM
151670
Manual annotation
SCANPY: ARI=0.34
BayesSpace: ARI=0.43
SpaGCN: ARI=0.36
SEDR: ARI=0.32
STAGATE: ARI=0.38
DeepST: ARI=0.35
GraphST: ARI=0.58
stGCL: ARI=0.44
Layer 3
Layer 4
Layer 5
Layer 6
WM
151671
Manual annotation
SCANPY: ARI=0.41
BayesSpace: ARI=0.48
SpaGCN: ARI=0.54
SEDR: ARI=0.55
STAGATE: ARI=0.6
DeepST: ARI=0.48
GraphST: ARI=0.63
stGCL: ARI=0.71
Layer 3
Layer 4
Layer 5
Layer 6
WM
151672
Manual annotation
SCANPY: ARI=0.38
BayesSpace: ARI=0.43
SpaGCN: ARI=0.57
SEDR: ARI=0.35
STAGATE: ARI=0.57
DeepST: ARI=0.5
GraphST: ARI=0.61
stGCL: ARI=0.76
Layer 3
Layer 4
Layer 5
Layer 6
WM
151673
Manual annotation
SCANPY: ARI=0.26
BayesSpace: ARI=0.56
SpaGCN: ARI=0.46
SEDR: ARI=0.49
STAGATE: ARI=0.5
DeepST: ARI=0.54
GraphST: ARI=0.62
stGCL: ARI=0.5
Layer 1
Layer 2
Layer 3
Layer 4
Layer 5
Layer 6
WM
151674
Manual annotation
SCANPY: ARI=0.35
BayesSpace: ARI=0.3
SpaGCN: ARI=0.39
SEDR: ARI=0.5
STAGATE: ARI=0.63
DeepST: ARI=0.38
GraphST: ARI=0.49
stGCL: ARI=0.66
Layer 1
Layer 2
Layer 3
Layer 4
Layer 5
Layer 6
WM
151675
Manual annotation
SCANPY: ARI=0.2
BayesSpace: ARI=0.35
SpaGCN: ARI=0.45
SEDR: ARI=0.52
STAGATE: ARI=0.45
DeepST: ARI=0.41
GraphST: ARI=0.62
stGCL: ARI=0.52
Layer 1
Layer 2
Layer 3
Layer 4
Layer 5
Layer 6
WM
151676
Manual annotation
SCANPY: ARI=0.19
BayesSpace: ARI=0.36
SpaGCN: ARI=0.31
SEDR: ARI=0.53
STAGATE: ARI=0.6
DeepST: ARI=0.39
GraphST: ARI=0.57
stGCL: ARI=0.55
Layer 1
Layer 2
Layer 3
Layer 4
Layer 5
Layer 6
WM

**Fig. S1.** **Comparison of spatial domains identified by stGCL, GraphST, DeepST, STAGATE, SEDR, SpaGCN, BayesSpace, SCANPY, and manually annotated layer structures in 12 slices of the DLPFC dataset.**

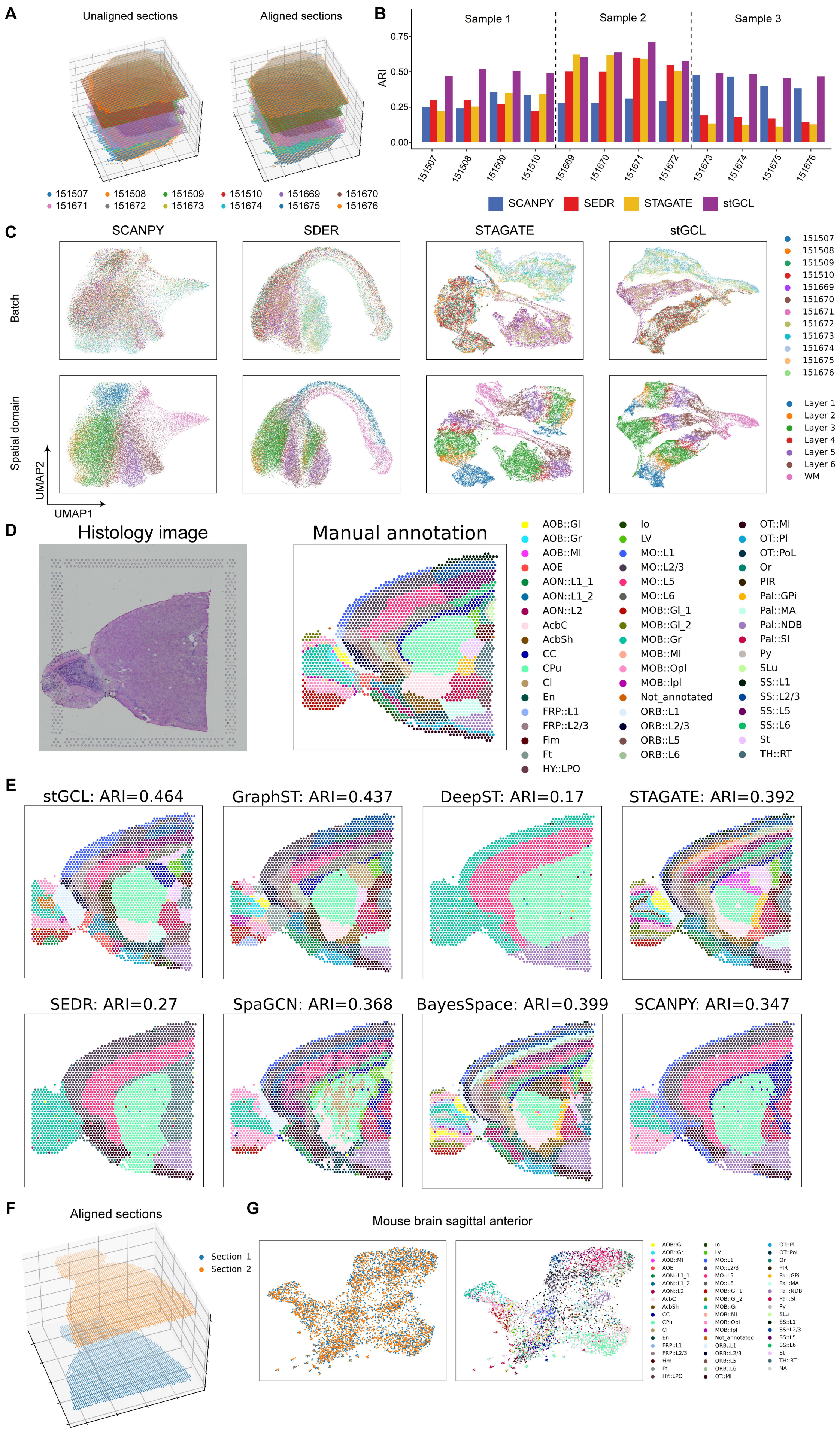

A
Unaligned sections
Aligned sections
151507 151508 151509 151510 151669 151670
151671 151672 151673 151674 151675 151676
B
Sample 1
Sample 2
Sample 3
ARI
SCANPY SEDR STAGATE stGCL
C
SCANPY
SDER
STAGATE
stGCL
Batch
Spatial domain
UMAP1
UMAP2
Layer 1 Layer 2 Layer 3 Layer 4 Layer 5 Layer 6 WM
D
Histology image
Manual annotation
E
stGCL: ARI=0.464
GraphST: ARI=0.437
DeepST: ARI=0.17
STAGATE: ARI=0.392
SEDR: ARI=0.27
SpaGCN: ARI=0.368
BayesSpace: ARI=0.399
SCANPY: ARI=0.347
F
Aligned sections
Section 1
Section 2
G
Mouse brain sagittal anterior

**Fig. S2. Results of integration of multiple tissue sections.** (**A**) Visualization of 12 DLPFC slices in 3D space (left); stGCL vertically aligns 12 DLPFC slices in 3D space (right). (**B**) Histogram of the spatial domain clustering results of the four methods across 12 tissue slices, in terms of ARI. (**C**) UMAP plots of 4 method embeddings on 12 slices of three samples. The spots are colored by slices (top) and cortical layers (bottom), respectively. (**D**) H&E image and manual annotation for the mouse brain sagittal anterior of section 1. (**E**) Spatial domains detected by different methods (including stGCL, GraphST, DeepST, STAGATE, SEDR, SpaGCN, BayesSpace and SCANPY algorithms). (**F**) stGCL vertically aligns 2 mouse brain sagittal anterior samples in 3D space. (**G**) UMAP plots of stGCL embeddings colored by the sections (left) and identified spatial domains (right) for integration of 2 mouse brain anterior samples.

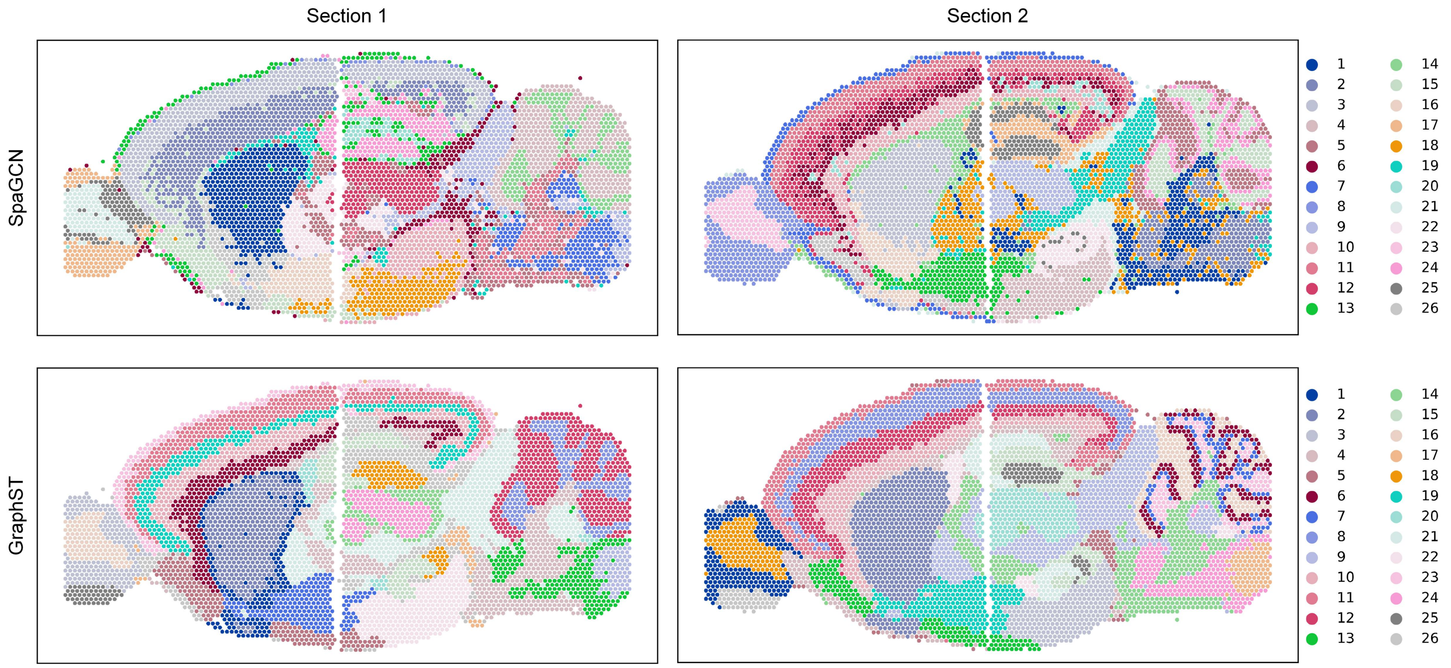


**Fig. S3. Results of horizontal integration of SpaGCN and GraphST on two mouse brain sections.**

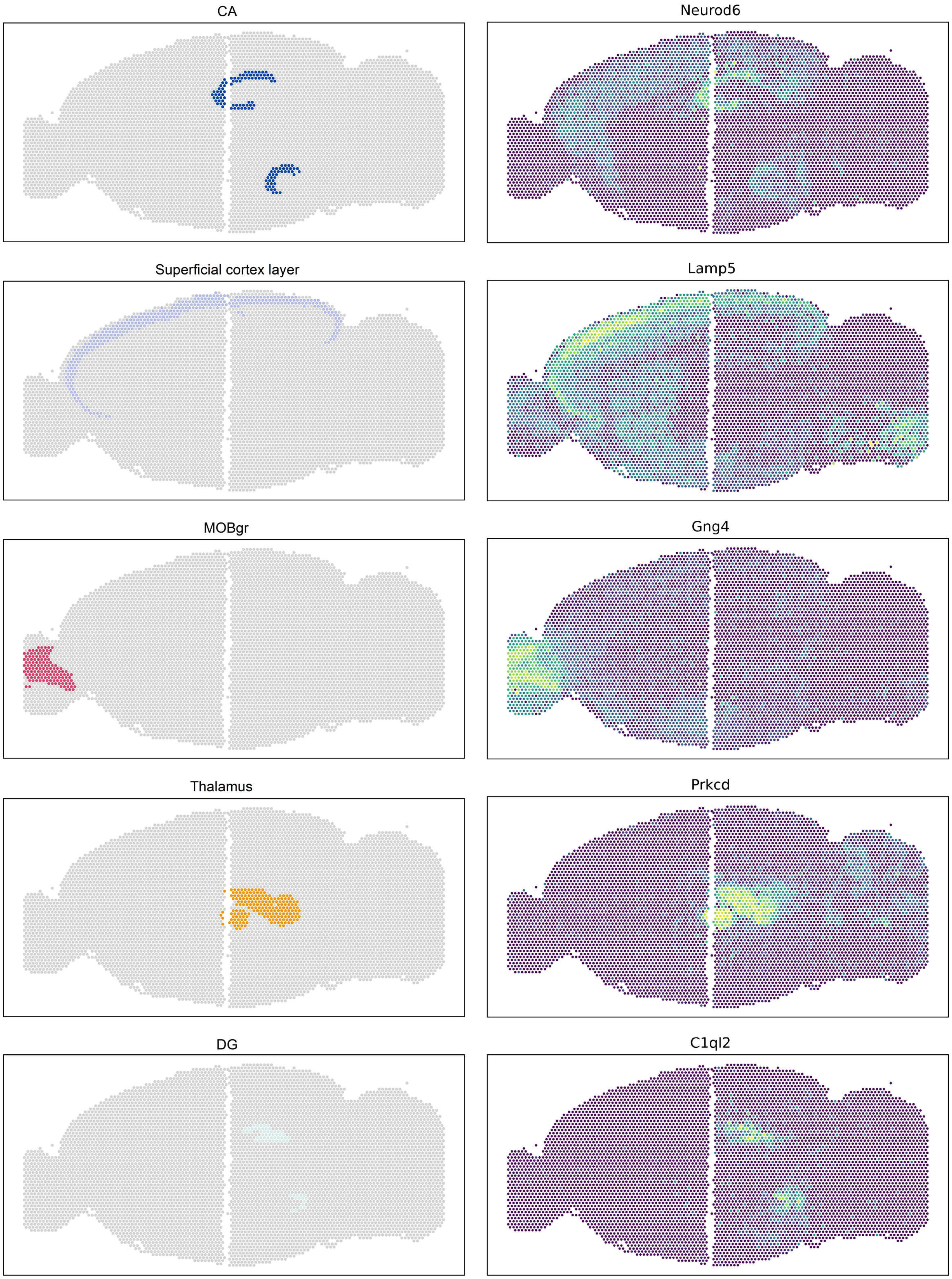


**Fig. S4. Visualization of the spatial domains (left) identified by stGCL and their corresponding marker genes (right).**

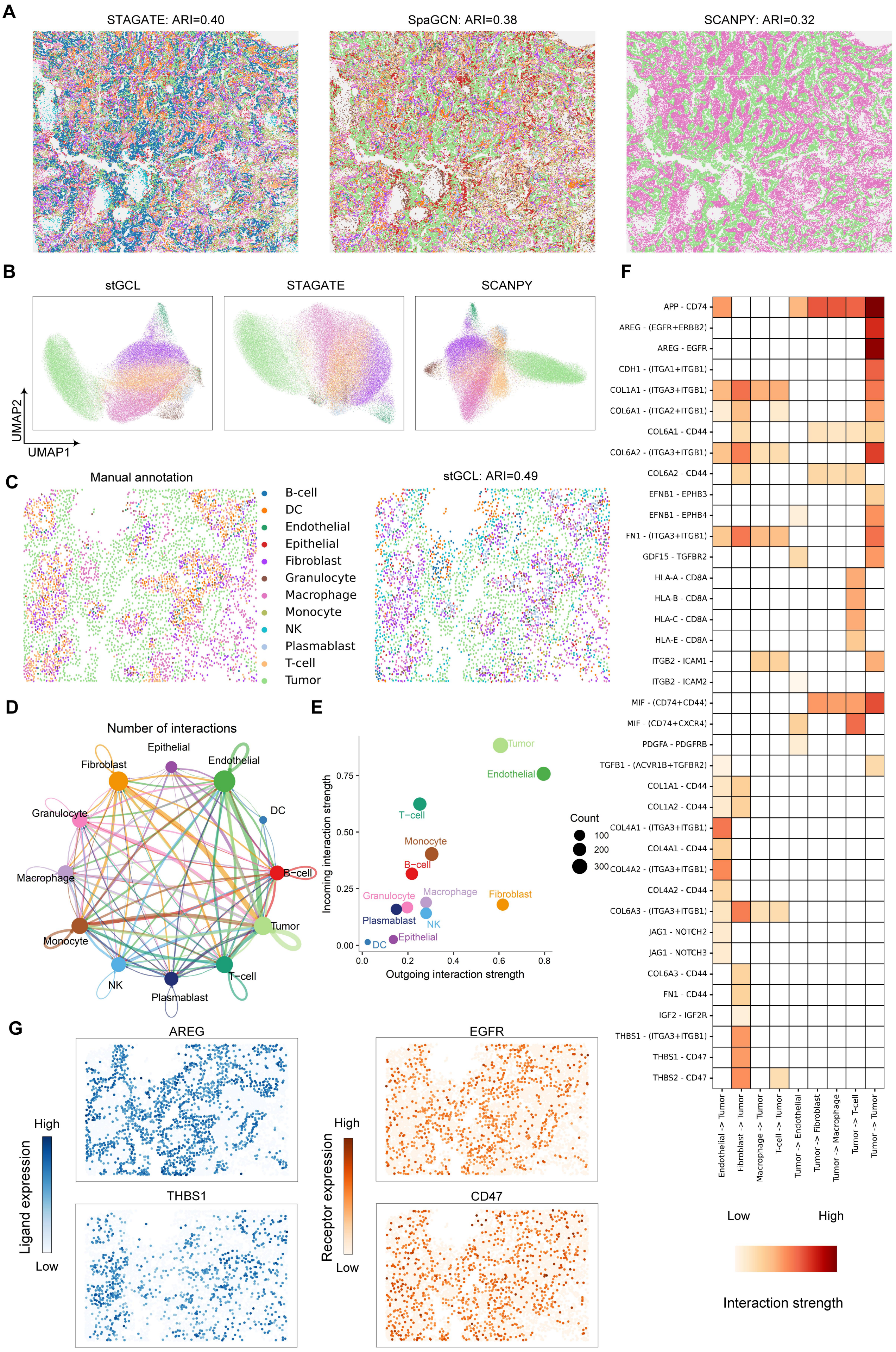
A
STAGATE: ARI=0.40
SpaGCN: ARI=0.38
SCANPY: ARI=0.32
B
stGCL
STAGATE
SCANPY
UMAP2
UMAP1
C
Manual annotation
B-cell
DC
Endothelial
Epithelial
Fibroblast
Granulocyte
Macrophage
Monocyte
NK
Plasmablast
T-cell
Tumor
stGCL: ARI=0.49
D
Number of interactions
E
Incoming interaction strength
Outgoing interaction strength
Count
100
200
300
F
APP - CD74
AREG - (EGFR+ERBB2)
AREG - EGFR
CDH1 - (ITGA1+ITGB1)
COL1A1 - (ITGA3+ITGB1)
COL6A1 - (ITGA2+ITGB1)
COL6A1 - CD44
COL6A2 - (ITGA3+ITGB1)
COL6A2 - CD44
EFNB1 - EPHB3
EFNB1 - EPHB4
FN1 - (ITGA3+ITGB1)
GDF15 - TGFBR2
HLA-A - CD8A
HLA-B - CD8A
HLA-C - CD8A
HLA-E - CD8A
ITGB2 - ICAM1
ITGB2 - ICAM2
MIF - (CD74+CD44)
MIF - (CD74+CXCR4)
PDGFA - PDGFRB
TGFB1 - (ACVR1B+TGFBR2)
COL1A1 - CD44
COL1A2 - CD44
COL4A1 - (ITGA3+ITGB1)
COL4A1 - CD44
COL4A2 - (ITGA3+ITGB1)
COL4A2 - CD44
COL6A3 - (ITGA3+ITGB1)
JAG1 - NOTCH2
JAG1 - NOTCH3
COL6A3 - CD44
FN1 - CD44
IGF2 - IGF2R
THBS1 - (ITGA3+ITGB1)
THBS1 - CD47
THBS2 - CD47
Endothelial -> Tumor
Fibroblast -> Tumor
Macrophage -> Tumor
T-cell -> Tumor
Tumor -> Endothelial
Tumor -> Fibroblast
Tumor -> Macrophage
Tumor -> T-cell
Tumor -> Tumor
Low
High
Interaction strength
G
AREG
THBS1
EGFR
CD47
Ligand expression
Receptor expression
High
Low

**Fig. S5. Comparisons of clustering results from different methods, and stGCL reveals L-R interactions in NSCLC data profiled by NanoString CosMx SMI.** (**A**) Clustering results obtained by STAGATE, SpaGCN and SCANPY at 20 FOVs of NSCLC data. (**B**) UMAP visualization produced by stGCL, STAGATE and SCANPY. The spots are colored based on the cell types identified by each method. (**C**) The manual annotations of NSCLC from FOV 2 and the cluster assignments generated by stGCL. (**D**) The cell-cell communication network identified on FOV 2 using cellchat based on stGCL embeddings. The edge width is proportional to the number of L-R interactions. (**E**) The scatter plot visualizes the senders (ligands) and receivers (receptors) in a 2D space. The x-axis and y-axis represent the interaction strength involved for each cell type. (**F**) Heatmap depicting the strength of L-R interactions between identified major cell types. (**G**) Spatial expression map of identified L-R pairs (AREG-EGFR, THBS1-CD47).

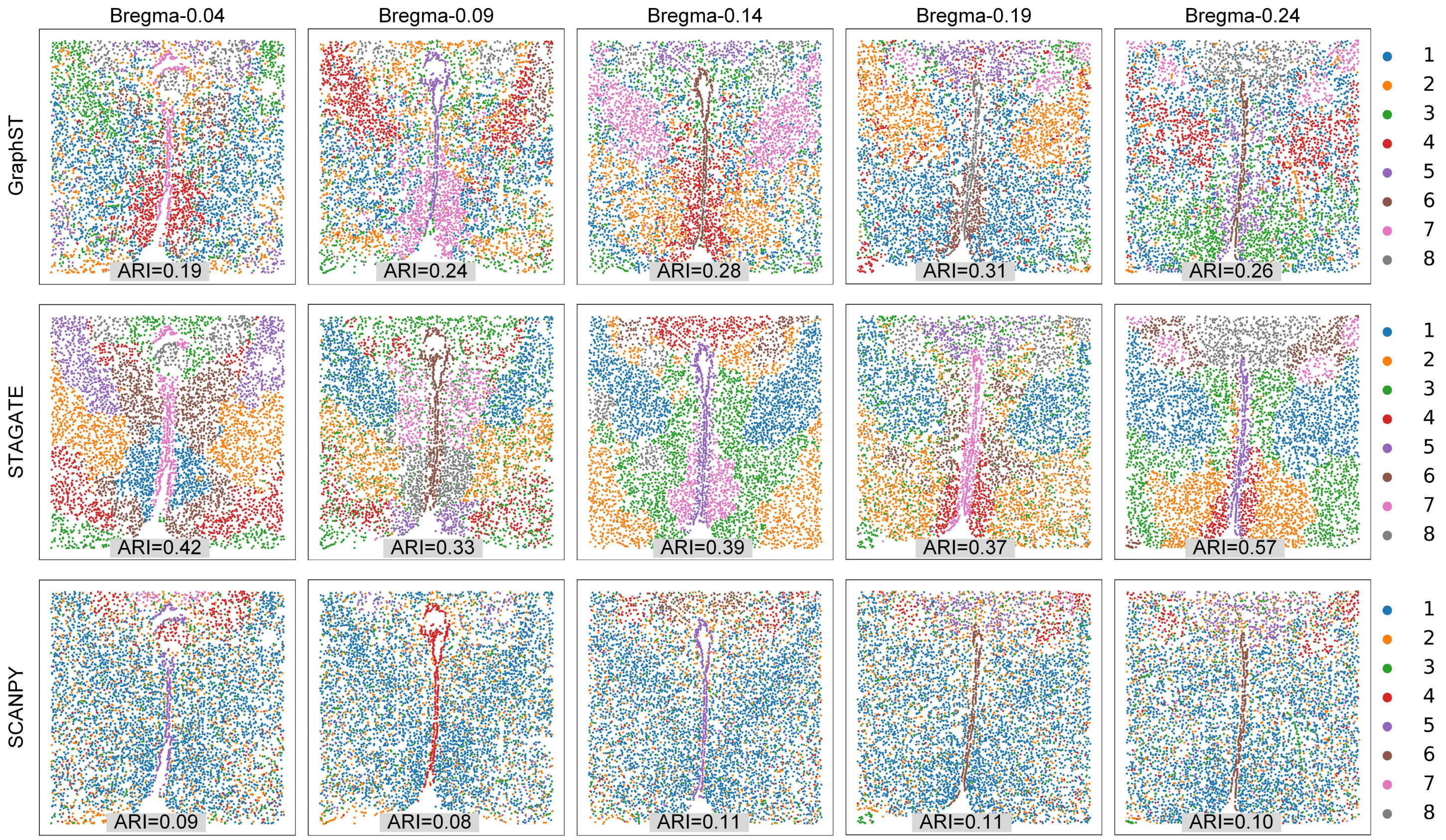


**Fig. S6. Clustering results obtained by GraphST, STAGATE and SCANPY methods on the mouse hypothalamic preoptic area MERFISH dataset.**

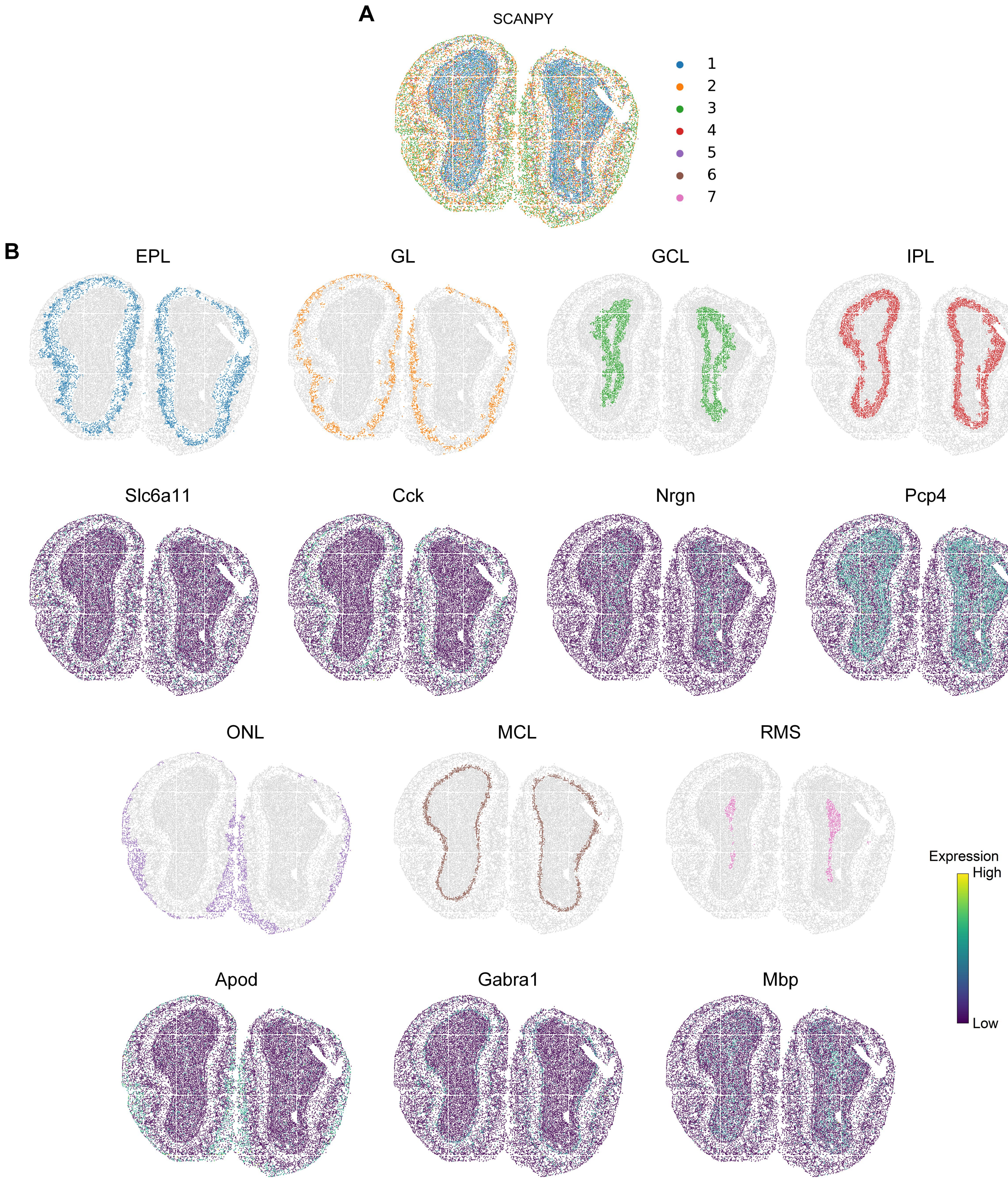


**Fig. S7. Spatial clustering on the mouse olfactory bulb Stereo-seq data.** (**A**) The spatial domain identified by SCANPY. (**B**) Visualization of the laminar structures detected by GraphST embeddings and the corresponding marker gene expressions.

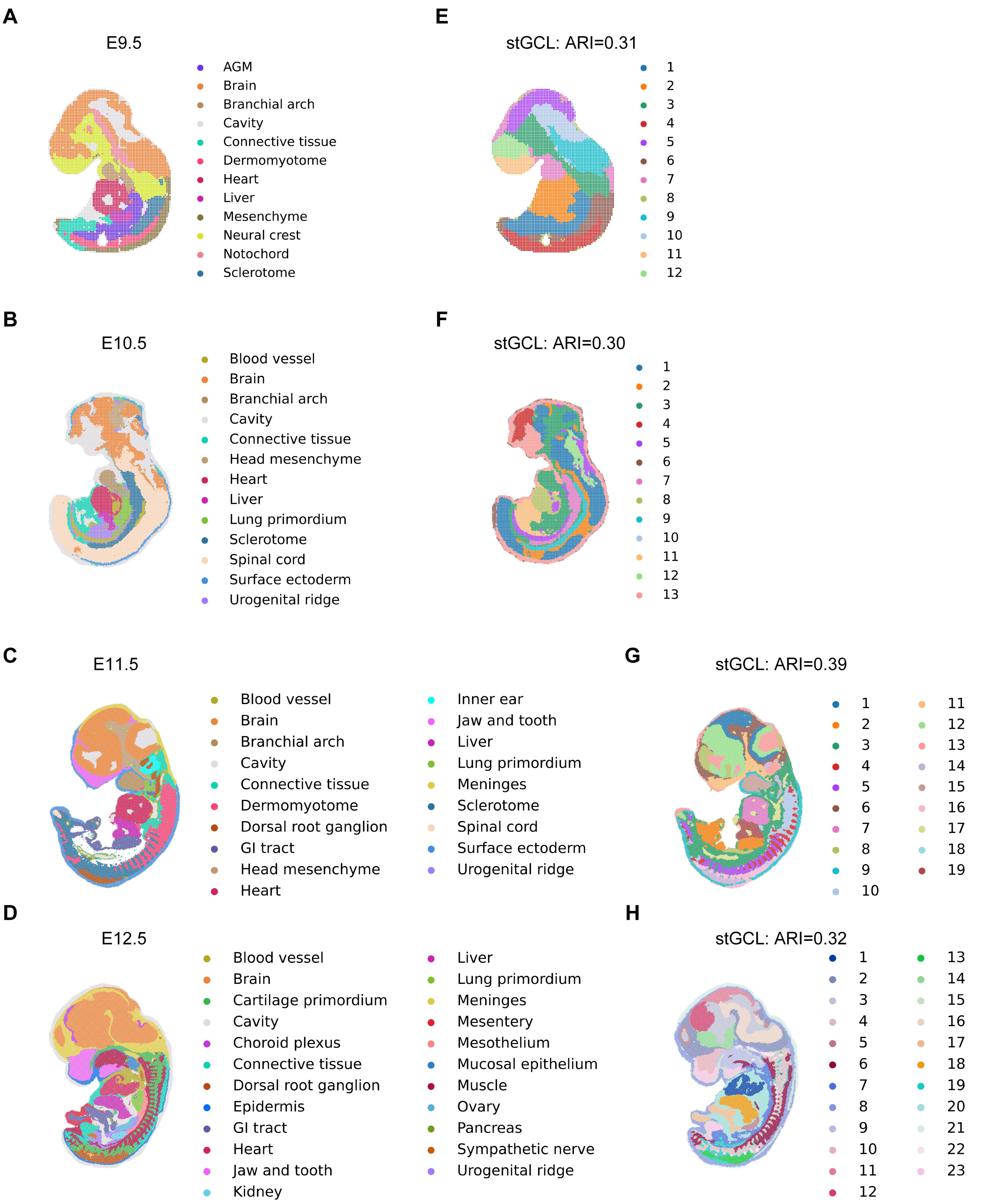


**Fig. S8. stGCL work on four Stereo-seq datasets.** (**A**), (**B**), (**C**) and **(D)** Manual annotations in the mouse embryos Stereo-seq datasets (E9.5, E10.5, E11.5 and E12.5). (**E**), (**F**), (**G**) and **(H)** Spatial domain identification is performed using stGCL and clustering results are measured by ARI.

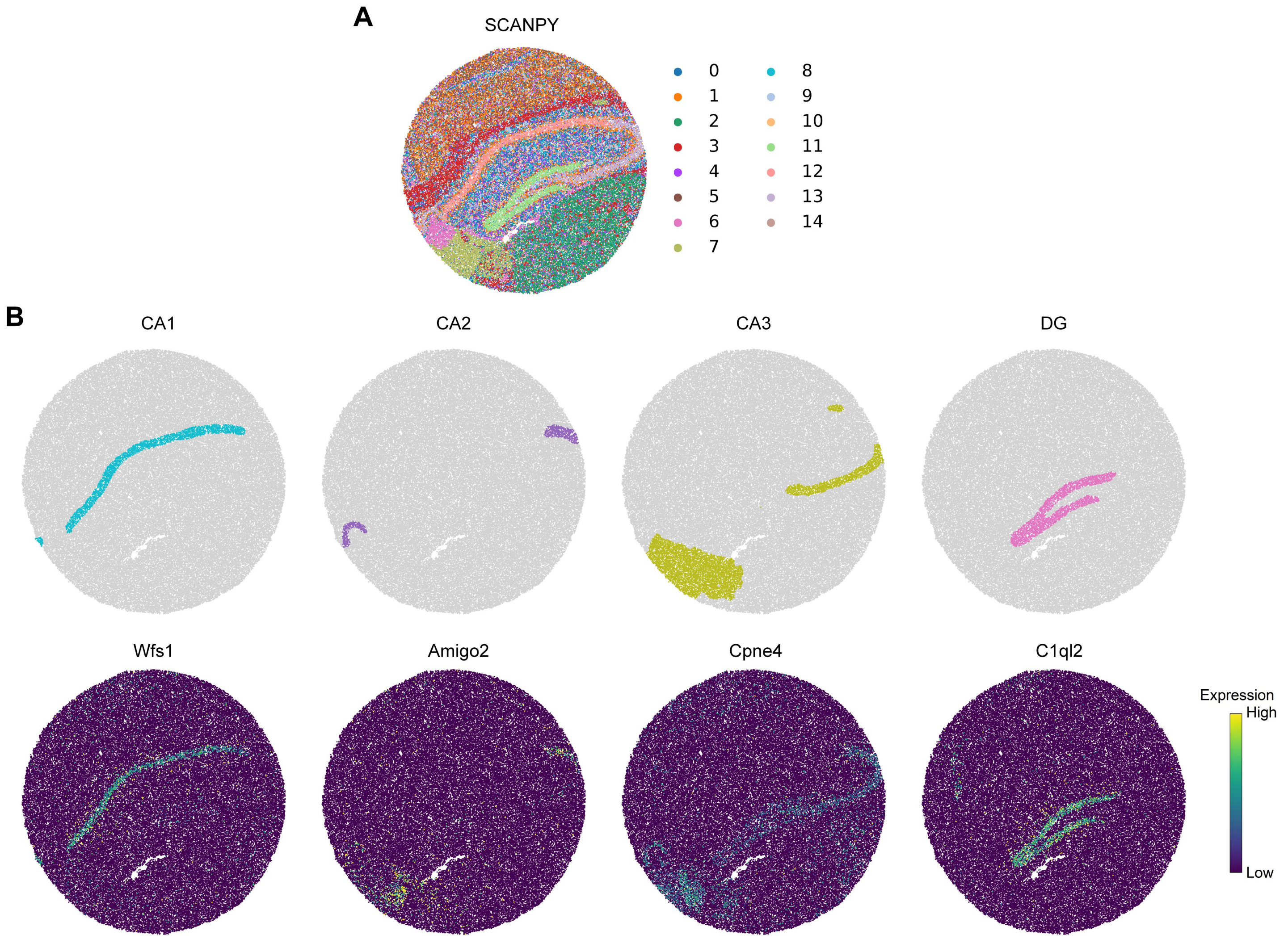


**Fig. S9. Spatial clustering on the mouse hippocampus Slide-seqV2 dataset.** (**A**) Spatial domains generated by SCANPY. (**B**) Visualization of the CA1, CA2, CA3, and DG structures identified by stGCL and their corresponding marker genes.

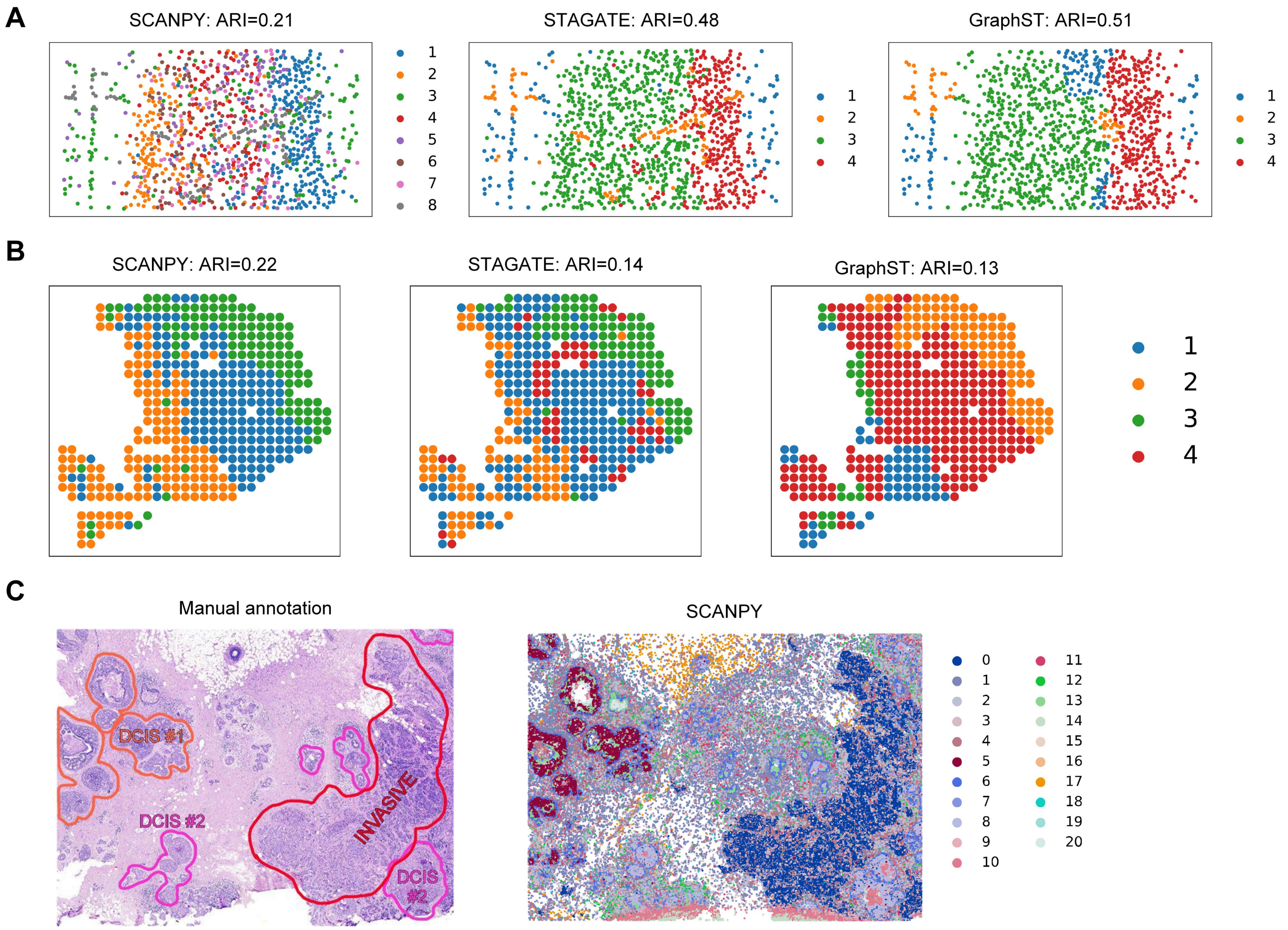


**Fig. S10. Detecting spatial domains of ST data from different platforms.** (**A**) Visualization of the spatial domain identified by different methods (GraphST, STAGATE and SCANPY) on the mouse mPFC STARmap dataset. (**B**) Clustering results using GraphST, STAGATE and SCANPY on PDAC data profiled by ST platform. (**C**) Manual annotations of breast cancer 10x Xenium data from the original study, and spatial domains detected by SCANPY.

A

| Tissue | Platform | Spots | Edges | Available histology images | Runtime (s) | GPU memory (MB) |
|---|---|---|---|---|---|---|
| DLPFC 151674 | 10x Visium | 3,673 | 21,258 | √ | 54.9 | 2015 |
| Mouse visual cortex | STARmap | 1,207 | 7,838 | × | 29.5 | 1343 |
| Mouse olfactory bulb | Stereo-seq | 19,109 | 291,130 | × | 249.6 | 3575 |
| Mouse hippocampus | Slide-seqV2 | 52,869 | 742,736 | × | 690 | 7805 |
| Breast cancer | 10x Xenium | 167,780 | 1373,536 | √ | 841.5 | 26869 |

B

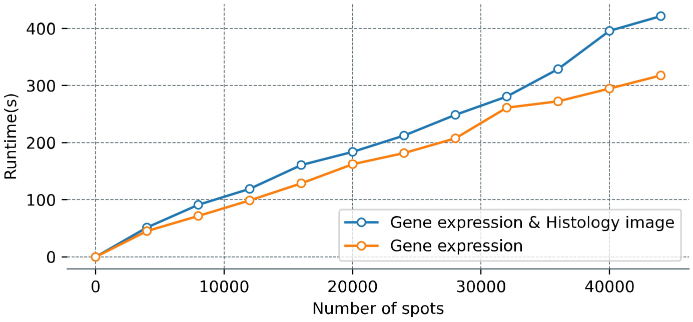


C

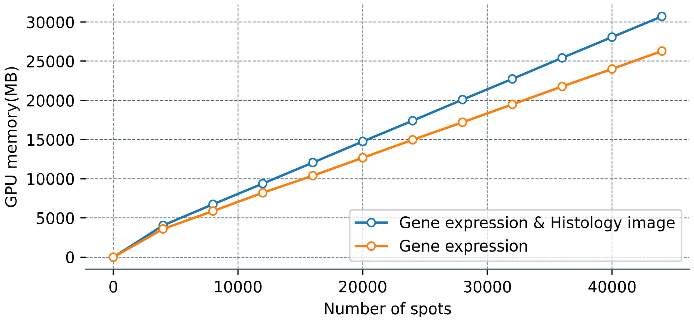


**Fig. S11. The computational cost of stGCL.** (**A**) The runtime and GPU memory usage of stGCL on five real datasets. (**B**) The runtime of stGCL on simulated datasets. (**C**) The GPU memory usage of stGCL on simulated datasets. The simulated datasets were obtained by replicating DLPFC slice 151674, one of which contained only gene expression modality data and the other consisted of gene expression modality and histological modality data. Experiments are performed on a PC with Intel(R) Xeon(R) Gold 6258R CPU @ 2.70GHz and NVIDIA QuADro GV100 GPU.

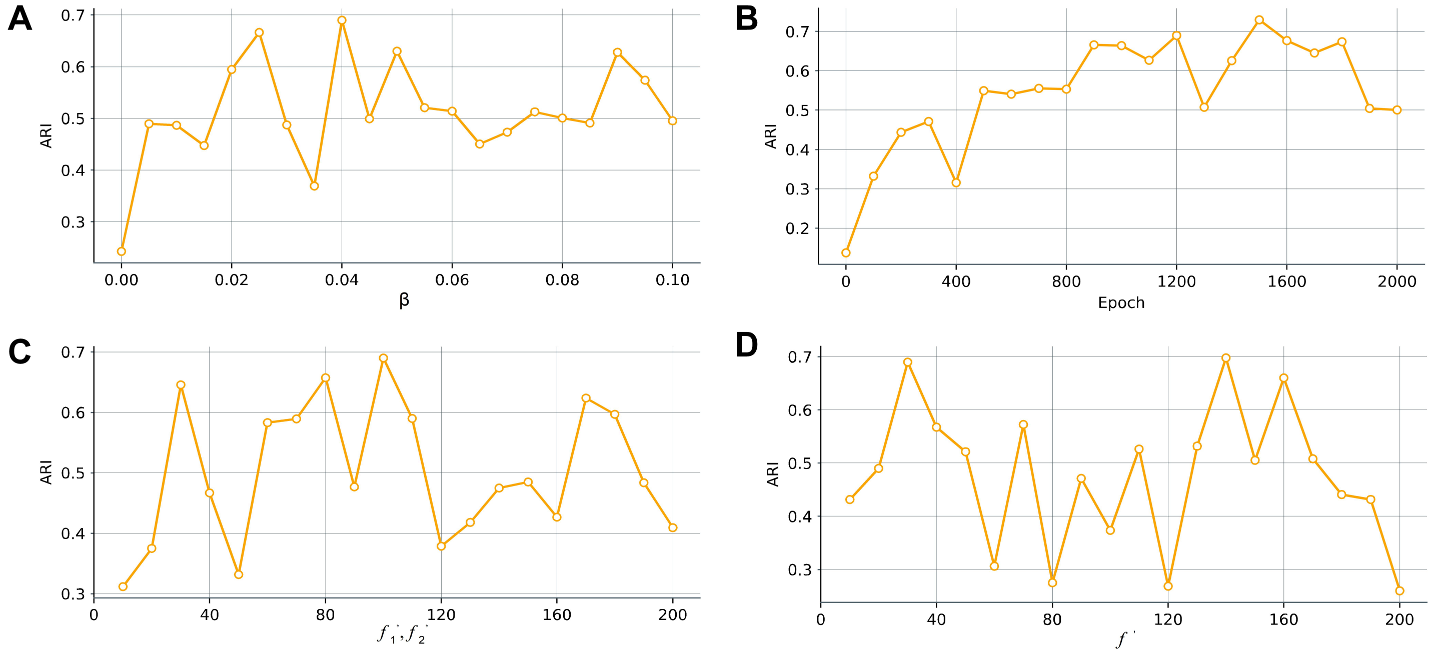


**Fig. S12. Clustering accuracy of stGCL at different hyperparameters selected by grid search on DLPFC slice 151674.** (**A**) Clustering results of stGCL under different β values. (**B**) Clustering results of stGCL at different epoch numbers. (**C**) and **(D)** Clustering results of stGCL under different latent dimensions of multi-modal GATE.

**Table S1. Summary of the ST data used in this study.**

| Tissue | Platform | Slice id | # of spots | Related figures |
|---|---|---|---|---|
| **Human dorsolateral prefrontal cortex (DLPFC)** | 10x Visium | 151507, 151508, 151509, 151510, 151669, 151670, 151671, 151672, 151673, 151674, 151675, 151676. | 4226, 4384, 4789, 4634, 3661, 3498, 4110, 4015, 3639, 3673, 3592, 3460. | Figure 2,3 Figure S1, S2 |
| **Mouse brain** | 10x Visium | Mouse Brain Section 1 (Sagittal-anterior) | 2695, 3355, 2825, 3289. | Figure 3 Figure S2, S3, S4 |
| | | Mouse Brain Section 1 (Sagittal-posterior) | | |
| | | Mouse Brain Section 2 (Sagittal-anterior) | | |
| | | Mouse Brain Section 2 (Sagittal-posterior) | | |
| **Mouse olfactory bulb** | Stereo-seq | N.A. | 19109 | Figure 4 Figure S7 |
| **Mouse embryo** | Stereo-seq | E9.5 | 5913, 18408, 30124, 51365. | Figure S8 |
| | | E10.5 | | |
| | | E11.5 | | |
| | | E12.5 | | |
| **Mouse hippocampus** | Slide-seqV2 | Puck_200115_08 | 52869 | Figure 4 Figure S9 |
| **Bronchiolar adenoma (BA)** | 10x Visium | N.A. | 4002 | Figure 5 |
| **Mouse medial prefrontal cortex (mPFC)** | STARmap | 20180419_BZ9_control | 1053 | Figure 4 Figure S10 |
| **Mouse hypothalamus** | MERFISH | Bregma-0.04, Bregma-0.09, Bregma-0.14, Bregma-0.19, Bregma-0.24 | 5488, 5557, 5926 5803, 5543, | Figure 4 Figure S6 |
| **Human breast cancer** | 10x Xenium | N.A. | 167780 | Figure 4 Figure S10 |
| **Pancreatic cancer** | ST | N.A. | 428 | Figure 4 Figure S10 |
| **Non-small cell lung cancer** | NanoString CosMx SMI | N.A. | 82843 | Figure 4 Figure S5 |

**Table S2. Sample ID and URL of anatomical reference data downloaded from the Allen Mouse Brain Atlas in this study.**

| Related figures | ABA ID | ABA URL |
|---|---|---|
| Figure 3E | 100883818 | http://atlas.brain-map.org/atlas?atlas=2&plate=100883818 |
| Figure 4H | 100960084 | http://atlas.brain-map.org/atlas?atlas=1&plate=100960084 |
